# Towards a digital diatom: image processing and deep learning analysis of *Bacillaria paradoxa* dynamic morphology

**DOI:** 10.1101/2019.12.21.885897

**Authors:** Bradly Alicea, Richard Gordon, Thomas Harbich, Ujjwal Singh, Asmit Singh, Vinay Varma

## Abstract

Recent years have witnessed a convergence of data and methods that allow us to approximate the shape, size, and functional attributes of biological organisms. This is not only limited to traditional model species: given the ability to culture and visualize a specific organism, we can capture both its structural and functional attributes. We present a quantitative model for the colonial diatom *Bacillaria paradoxa*, an organism that presents a number of unique attributes in terms of form and function. To acquire a digital model of *B. paradoxa*, we extract a series of quantitative parameters from microscopy videos from both primary and secondary sources. These data are then analyzed using a variety of techniques, including two rival deep learning approaches. We provide an overview of neural networks for non-specialists as well as present a series of analysis on *Bacillaria* phenotype data. The application of deep learning networks allows for two analytical purposes. Application of the DeepLabv3 pre-trained model extracts phenotypic parameters describing the shape of cells constituting *Bacillaria* colonies. Application of a semantic model trained on nematode embryogenesis data (OpenDevoCell) provides a means to analyze masked images of potential intracellular features. We also advance the analysis of *Bacillaria* colony movement dynamics by using templating techniques and biomechanical analysis to better understand the movement of individual cells relative to an entire colony. The broader implications of these results are presented, with an eye towards future applications to both hypothesis-driven studies and theoretical advancements in understanding the dynamic morphology of *Bacillaria*.

## Introduction

“I still remember, as many years ago, when I found the *Bacillaria* paradoxa near Greifswald many years ago, that I stood as if clinging to the microscope and could not turn my back on the strange spectacle that presented itself to me…. they are glued together as if they were an organism, and yet each moves for itself next to the other!”, translated from Max Schultze [1].

Creating digital instantiations of a model organism is of great potential to well-established communities centered around model organisms such as *Caenorhabditis elegans* [2]. The opportunity for creating a digital model of a non-model organism is potentially greater. In this paper, we will introduce a data-intensive approach to modeling the behaviors and dynamic phenotypes associated with the colonial diatom *Bacillaria paradoxa* (Figure 1). Using image processing techniques, we can construct a computational model of *Bacillaria* colonies. This simple phenotypic model provides a means to capture the dynamics of movement across different moments in time. Inferring the movement of these colonies reveals the diversity and complexity of behavior exhibited in a simple organismal colony.

**Figure 1.**
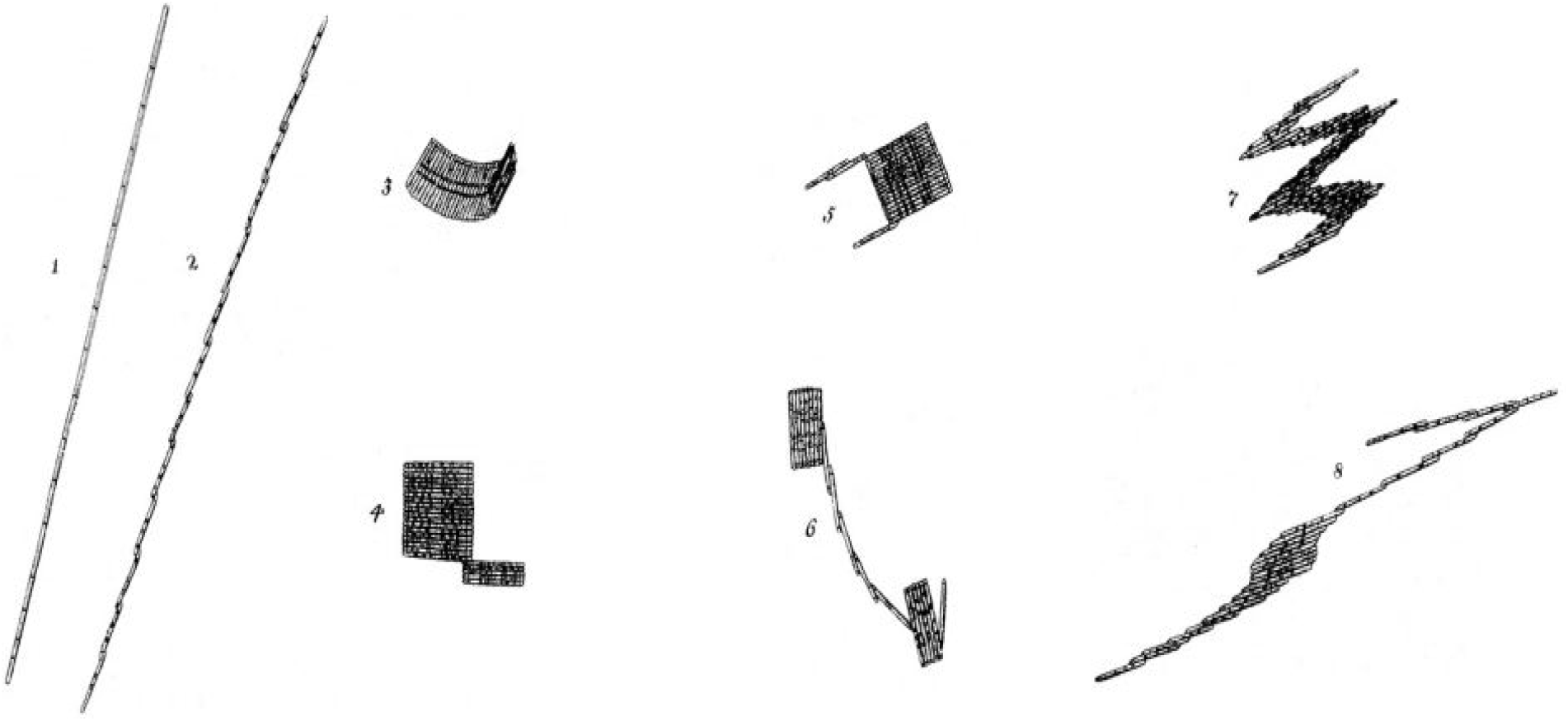
Drawing adapted from O.F. Müller (1783, translated in [3]), who was the first to characterize *Bacillaria* colonies. Examples 1 through 8 show the various states of expansion and contraction (dynamic phenotypes) of colonies.

The value of this approach is enhanced by the nature of the *Bacillaria* literature. While the literature is quite old (*Bacillaria* was first observed by Otto Friedric Müller in 1783 [3, 4], work to date has focused mostly on taxonomy and cellular/structural biology. The partial synchrony of *Bacillaria* colonies [5] indicates that the behavior of a colony may be greater than the sum of its individual cells (i.e. that each colony is a multicellular organism). Cellular biology work that has been done on the movement of *Bacillaria* [6–8], is neither computational in nature nor at the whole-organism level.

There has been some computational work conducted on diatom morphogenesis [9–18] which is expanding due to their usefulness in nanomaterials [19–23], medical applications [24], and many other fields [25]. Yet despite these studies, there is little computational work integrating structural morphology with diatom motility.

### Organism Description

*Bacillaria paradoxa*, synonymous with *Bacillaria paxillifera* [26], is a diatom in the *Bacillariaceae* family which has been subdivided into three species: *B. paxillifera, B. kuseliaeand* and *B. urve-millerae*, with *B. paxillifera* further divided into four varieties: var. *czarneckii*, var. *pacifica*, var. *tropica* and var. *tumidula* [26]. As the distinctions are mostly made at the SEM level of resolution, we will adopt the blanket designation *B. paradoxa* in this paper (Figure 1). Diatoms are a group of eukaryotic microalgae whose ornate cell walls are composed primarily of amorphous silica. They exhibit a unique life-history [27]. Cells of *Bacillaria* (sometimes called filaments) are elongated and motile, sliding along each other, in stacked colonies that curve slightly out of the plane. Cells are rectangular in girdle view (as seen in colonies, as their valves face each other), and lanceolate in valve view (Figure 2). The raphe system is slightly keeled and runs from pole to pole with no central nodule [28]. Two large plate-like chloroplasts are present, one near each end of the cell. The nucleus is located centrally. Cells are yellow-brown in color. Fibulae are strong, and the valve surface is covered in transverse parallel structures called striae [26].

**Figure 2.**
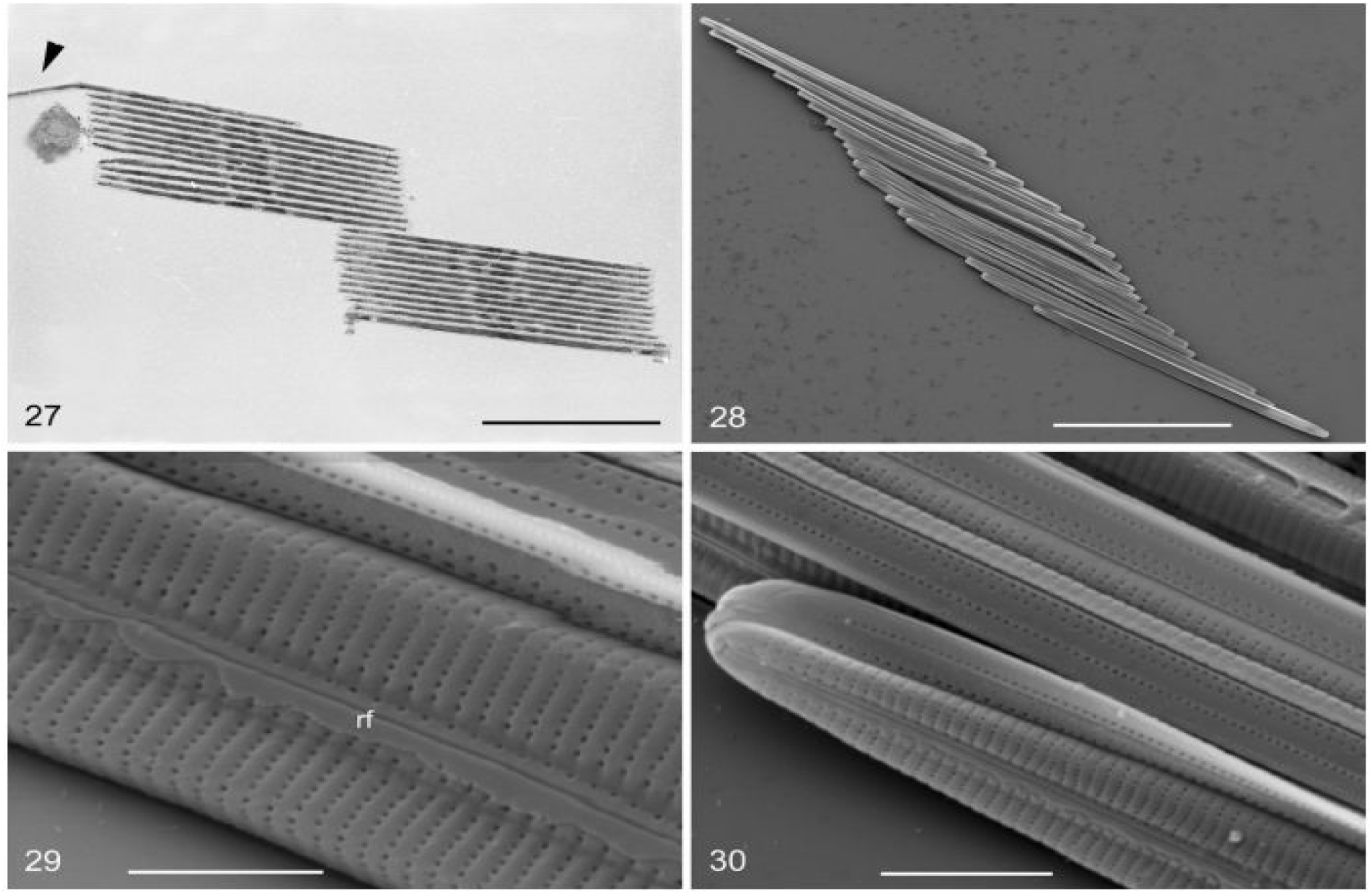
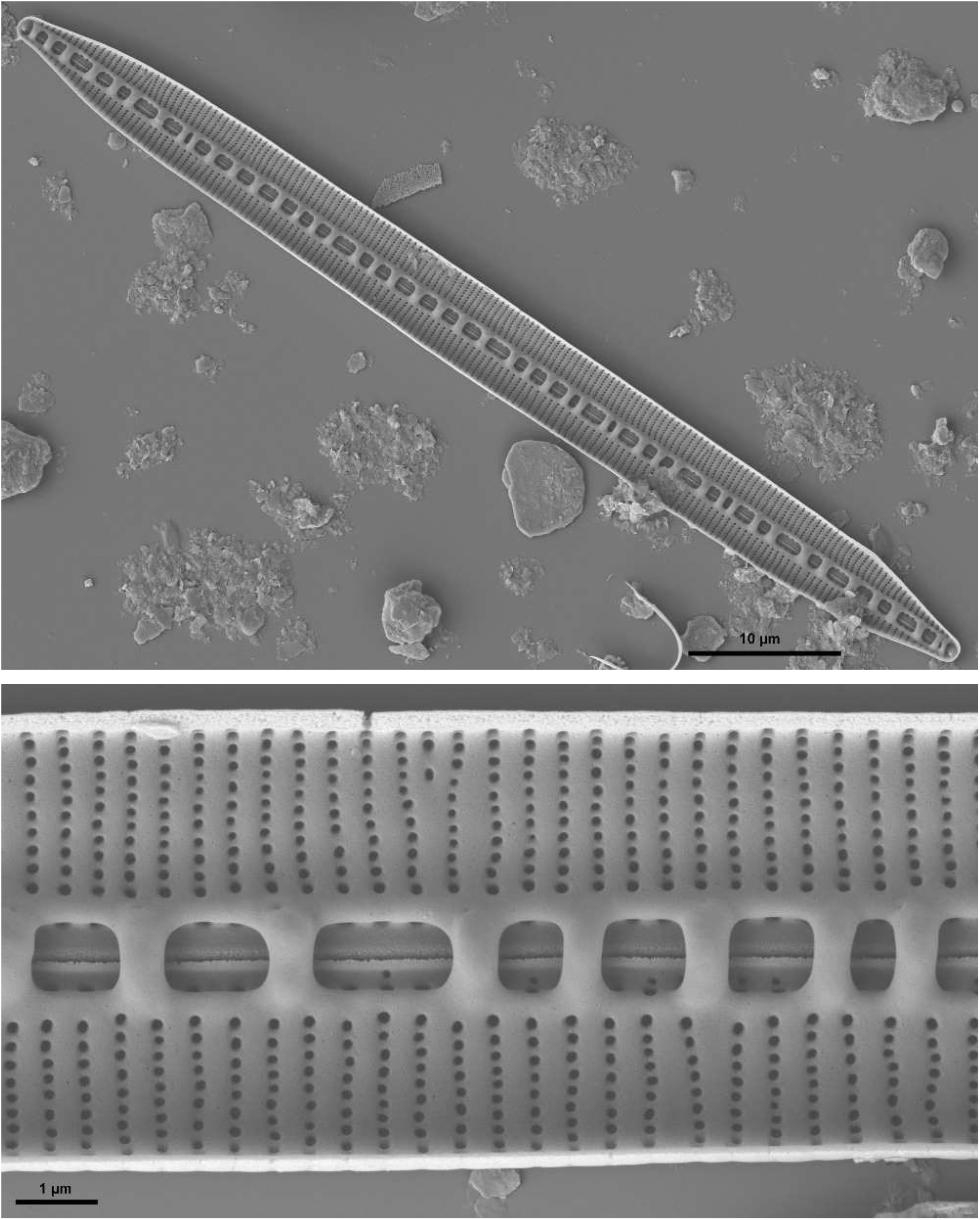
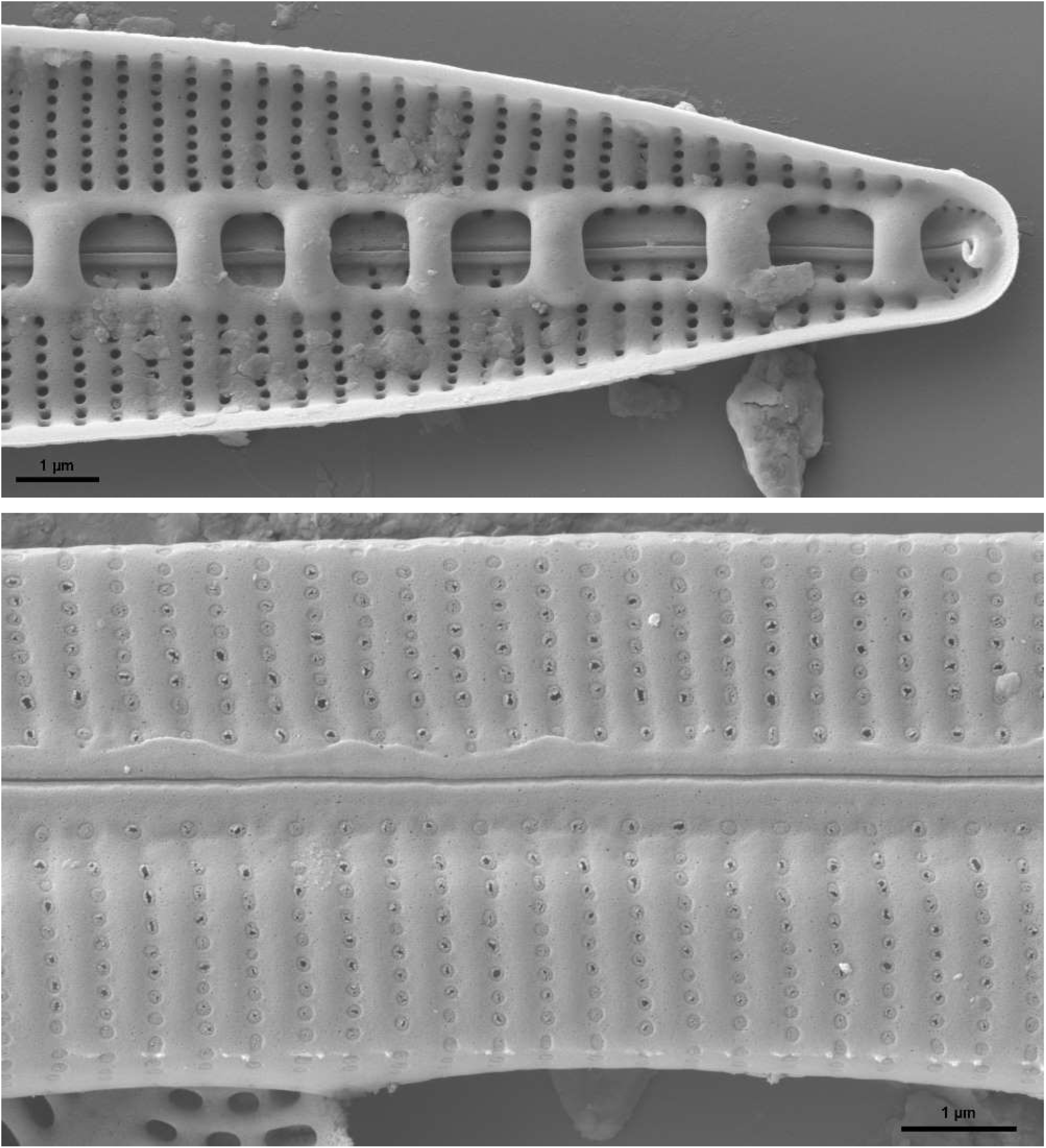
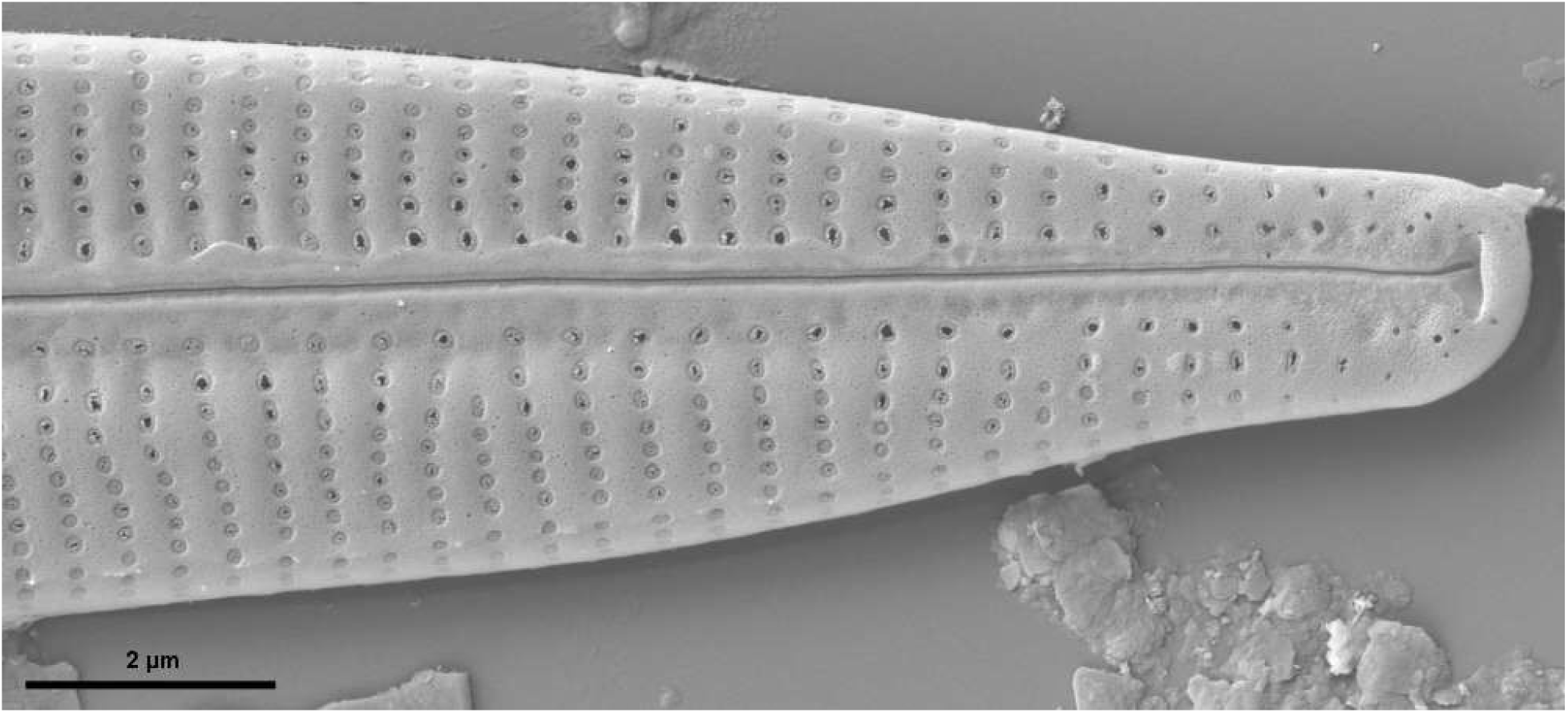
*Bacillaria* close-up images of single cells using Scanning Electron Microscopy (SEM). A: a whole valve sees from the inside. B: close up of the same, middle section. The horizontal slit is the raphe. It lacks a central node. C: Tip of the inside of a valve. D: Middle section of a valve, exterior view. Note the raphe is a slit through the whole valve. E: External view of the tip of a valve [37] with permission of Amgueddfa Cymru – National Museum Wales **[requested November, 20 2019]**.

*Bacillaria* cells are arranged in parallel stacks, and these parallel stacks form a colony. These stacks form early in the life-history of a colony by cell divisions perpendicular to the valves. The stacked colony moves by consecutive pairs of individual cells sliding against each other [3]. When these sliding movements occur in temporal order across the colony [5], the synchronized movement is like the sliding of a deck of cards [29] and results in a large extension of the whole colony.

The continual folding and unfolding of a colony of cells stacked in parallel results in cyclic gliding movements. Each cell is an intrinsic oscillator [30], so the partial synchronization may be due to entrainment of these oscillators by an unknown mechanism, perhaps light piping within each colony [31, 32]. The mechanism of gliding movement is still being worked out [33]. Observations of high accelerations of single diatoms [34] suggest a motor that moves with explosive force at a molecular scale [35]. This may be the basis of the often observed jerky motion of diatoms, hints of which have been seen in *Bacillaria* [36].

### Research Motivation

Diatoms (and *Bacillaria* in general) are an excellent model system for understanding phenotypic growth and dynamics [38]. Unfortunately, there is no model for its behavioral and dynamic phenotypic properties. It is our contention that digital technologies such as image processing techniques, machine learning techniques, and open data repositories allow us to construct a digital model that can be used to uncover previously unexplored relationships.

Our data rely on the ability to generate microscopy movies from cultured specimens. Some of these movies are publicly available in places such as YouTube or in the Supplemental Materials of academic papers. Fortunately, *Bacillaria* is relatively easy to isolate and culture, and we have collected primary microscopy data as well. Enabled through the application of deep learning [39–41] and biomechanical techniques, we postulate that a digital *Bacillaria* is not only possible but highly useful to the scientific community.

Our purpose here is twofold: to review key concepts related to our computational approach, and to present the technical details of building the digital *Bacillaria*. First, we will provide an overview of neural networks and machine intelligence. Then the methods used for extracting data from individual video images will be presented, along with more detailed descriptions for methods employed in the data analysis. The paper will conclude with an analysis using several techniques for image processing and discovering the computational features that define a *Bacillaria* colony. These include the shape parameters of the cell, techniques to extract intracellular features, and techniques to approximate colony movement. Our techniques range from formal deep learning techniques for extracting morphological features to the creation of masks and templates to quantify images extracted from time-series.

#### Description of Neural Networks

Neural Networks are computer programs assembled from many computational units that behave as an adaptive system. This adaptive system is inspired by the nervous system: providing the systematic power and computational capabilities of a computer with the densely reticulating connectivity of a model of a biological brain. The rationale for using neural networks (sometimes referred to a connectionist approach) in a computational setting is to simulate the pattern recognition and decision-making properties of biological learning.

Computers and brains have much in common, but there exist important differences. The most fundamental difference is that computers and minds think and reason (such as this exists in machines) in entirely different ways. Computational networks are also wired in very simple ways, usually in serial chains. Each link in this chain is connected to maybe four or five others in arrangements known as logic gates. By contrast, the brain has complex entities called neuronal cells which are densely interconnected in complex, parallel ways. Each neuron can be connected with up to 10,000 of its neighbors. These connection patterns lead to functional distinctions which are currently the subject of great debate [42].

Input units are designed to receive various forms of information from the outside world that the network will attempt to learn about, recognize, or otherwise process information. Other units reside on the opposite side of the network and signal how it responds to the information it’s learned; those are known as output units. Situated between input units and output units are one or more layers of hidden units, which, together, form the majority of the network. Most neural networks are fully connected, which means each hidden unit and each output unit is connected to every unit in the adjacent layers on either side. Connections between one unit and another are represented by a number called a “weight”, which can be either positive (if one unit excites another) or negative (if one unit suppresses or inhibits another). The higher the weight, the more influence one unit has on another. (This corresponds to the way actual brain cells trigger one another across tiny gaps called synapses, with excitatory or inhibitory connections.)

Neural networks learn things through training and testing. During both training (when the ANN learns associations) and testing (when the ANN makes associations), patterns of information are fed into the network via the input units. In turn, layers of hidden units are activated, which ultimately trigger the output units. This common design is called a feedforward network. Each unit receives inputs from the units to its left, and the inputs are multiplied by the weights of the connections they travel along (Figure 4). Every unit adds up all the inputs it receives. When the sum is more than a certain threshold value, the unit “fires” and triggers all connected units.

For a neural network to learn, there has to be an element of feedback involved—just as children learn by being told what they’re doing is right or wrong. In fact, we all use feedback, all the time. Neural networks learn things through a process of feedback called backpropagation (or backprop). This involves comparing the output a network produces with the output it was meant to produce, and then modify the weights of the connections between the units in the network accordingly. In time, backpropagation causes the network to adapt to the desired output (or learn), reducing the difference between actual and intended output to the point where the two coincide, so the network figures things out exactly as it should.

Once the network has been trained with enough learning examples, it reaches a point where you can present it with an entirely new set of inputs that it has never seen before and see how it responds. For example, suppose you’ve been teaching a network by showing it many pictures of chairs and tables, represented in some appropriate way it can understand, and telling it whether each one is a chair or a table. As an example, suppose we train the model with 25 images of chairs and 25 images of tables. Depending on how completely the model is trained, new examples will be categorized as either a chair or a table. This is generalized from its past experience rather than being generated *de novo*.

The inputs to a network are essentially binary numbers: each input unit is either switched on or switched off. Using five input units, we can feed in information about different features of an object using binary strings. For example, a chair might conform to five feature categories: possessing a back, possessing a top, having soft upholstery, allows for comfortable sitting, and storage capacity. This results in a series of categorical responses: Yes, No, Yes, Yes, No (10110). For a table, these responses might be: No, Yes, No, No, Yes (01001). During the learning phase, the network is simply looking at binary strings (e.g. 10110 and 01001), to determine what represents a chair and what represents a table.

## Methods

### Video extraction

Microscopy videos of *Bacillaria* were obtained from a number of sources (private archives and YouTube). The video frames were then processed to obtain segmented features and numeric variables. Specifically, we decomposed video files of *Bacillaria* microscopy images into its component frames. The number of frames depends on the sampling rate for the given video. Deep Learning models DeepLabv3 (a TensorFlow library reviewed in [43]) and OpenDevoCell (built with DeepLearning4J; see [44]) were then used to analyze these images. Due to the heterogeneous sources of acquisition, information was extracted in a manner that allows for unsupervised classification of the *Bacillaria* colony. The dataset for the DeepLabv3 analysis consisted of around 20,000 frames of secondary microscopy data, which contains much variety related to *Bacillaria* movement. The data for the OpenDevoCell and time-lapse analyses comes from microscopy data collected specifically for these analyses.

#### Image Skeleton Creation

The pre-masked images (skeletons) derived for our primary data were created in GIMP 2.10 [45]. First, we converted each raw image to an indexed (1-bit) imaged with no color dithering. Next, we converted the resulting binary map to an RGB (red, green, blue) indexed image. Select the cell area (RGB value 0,0,0) by color, and change to RGB value to (0,217,0). Maintaining the selection by color, edit the stroke selection function to a width of 1.0 pixels for a thin skeleton, and 5.0 pixels for a thick skeleton. This ensures separation between edges that are close together while also remaining selectable by the segmentation algorithm.

Once the boundaries (which were colored as 0,217,0) have been selected, select the area inside the boundary and change to RGB value 0,0,0. These transformations should result in a black cell surface area with a light green boundary. The final step is to change the background color (select the background by color) to RGB value 0,0,0. To create a thick skeleton from a thin skeleton, select the thin skeleton by color and then select the border function. The border width should be set to 4, hard border, and filled with RGB value 0,217,0. The pseudo-code for GIMP script-fu is located on Github (https://github.com/devoworm/Digital-Bacillaria/tree/master/Image-Skeletons).

#### Image Tracking for Movement

We also employ image tracking for the primary microscopy data. The tracking of a partial image (template) of a diatom can be used under certain conditions to obtain its trajectory. In particular, a movement of the diatoms in a plane perpendicular to the optical axis is essential. Sufficiently differentiated structures (chloroplasts) are required. The boundaries between neighboring diatoms are not considered.

Image tracking is done using Tracker 5.1.3 (https://www.softpedia.com/get/Science-CAD/Douglas-Tracker.shtml), an image analyzer that provides information regarding the motion of features between movie frames. For the tracking of paths in videos, various algorithms have been developed and implemented [46]. Each method has specific applications and strengths. It is often assumed that the objects to be tracked differ significantly from a sufficiently homogeneous background, yet this is not the case for *Bacillaria paxilifer* colonies.

Image tracking allows for cells in the colony to be tracked without complete separation from its background. Our ad-hoc method of feature selection defines cells as an ellipse registered with horizontal and vertical axes (Figure 3A). For sake of consistency, all video frames are rotated by 31 degrees and cropped. This was done primarily to make alignment of the ellipse and the diatom identical. This allows for templates of different size in addition to a direct comparison between cells using the x-axis. The tracking procedure proceeds by numbering the cells within a colony (see Figure 3B). The cell labeled “1” indicates the reference cell that is fixed relative to the substrate. Diatoms labeled “2” and “3” are then tracked as shown in the bottom panel of Figure 3. As these numeric values refer to the position of a particular cell relative to a reference cell (position of cell #2 relative to cell #1), an offset value must be added in order to optimize initial positions relative to movement.

**Figure 3.**
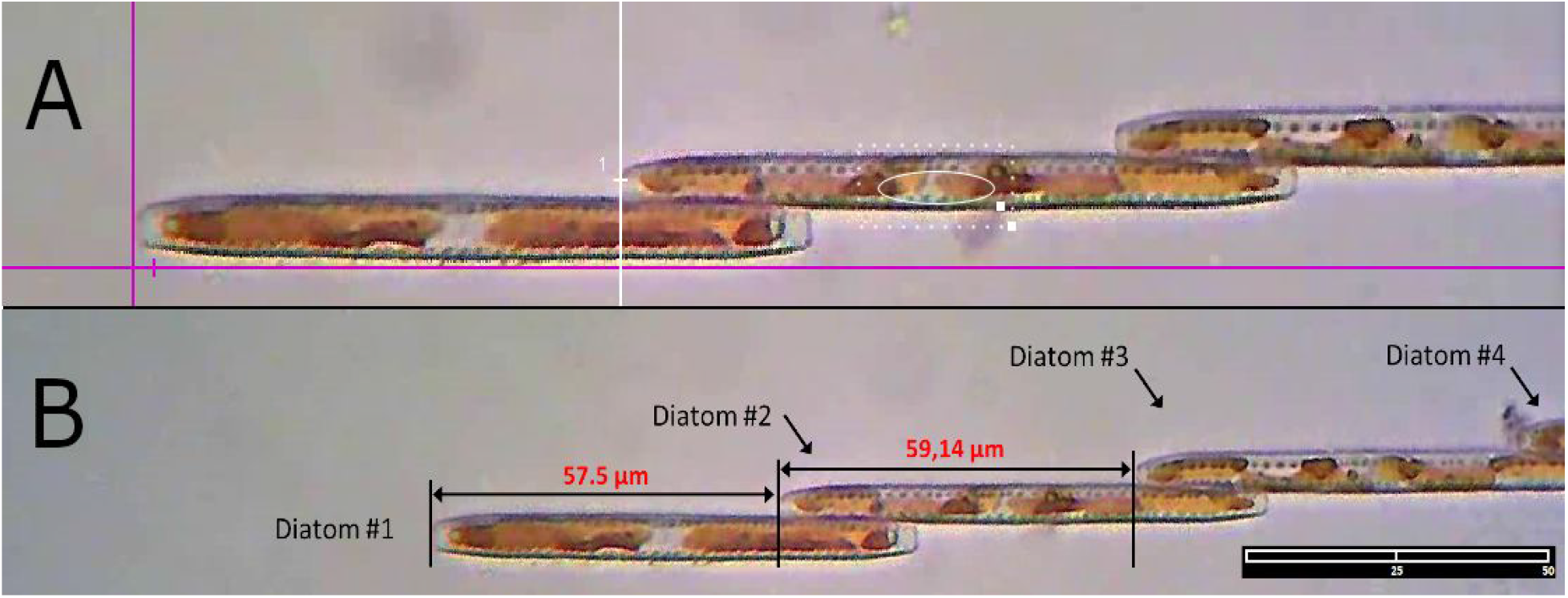
Demonstration of the image tracking procedure. A: definition of tracked feature (white ellipse within a cell). B: labeled (numbered) cells with relative measurements provided in red. The determined coordinates refer to a target in the middle of the template. The target can be moved and placed on the apex of the diatom being tracked. Then the coordinates of the apex are captured. This position is indicated in Figure 3A by a mark and a vertical line. Image scale: 38.36 μm per cm, or 0.325 μm per pixel. Scale bars 50 μm

### Deep Learning

Deep Learning [47, 48] is implemented using a pre-trained model called DeepLabv3 and open-source software with a web interface called OpenDevoCell (based on DeepLearning 4J). DeepLabv3 (Google, MountainView, California, USA) is a package for TensorFlow, and Deep Learning 4J (Eclipse Foundation, Ottawa, Canada), a Java-based library that works with TensorFlow. OpenDevoCell is open-source software located on Github (https://github.com/devoworm/GSOC-2019/tree/master/OpenDevoCell) and as a web-based application (https://open-devo-cell.herokuapp.com).

The Deeplab model [43] is implemented in TensorFlow, a platform for implementing Machine and Deep Learning algorithms. The Deeplab pre-trained model is based on deep learning, which utilizes a neural network of many layers. Deeplab is also specialized for extracting features that conform to a bounding box, as it has been trained on both “hard” object classes (objects embedded in an obscuring background) and datasets such as ImageNet and JFT-300M [49]. Bounding boxes of different sizes and orientations not only enable a heuristic means of segmenting individual cells in a *Bacillaria* colony, based on visual inspection it fits the cell shape and nature of variability quite well.

The Deeplab v3 algorithmic analysis is based on the idea of dilated convolution, where the input signal is sampled in alternative ways [49]. Ultimately, the algorithm attempts for find a tradeoff between localization (using a small sampling aperture of image pixels) and context assimilation (using a large sampling aperture of image pixels). This optimal tradeoff in spatial resolution determines which pixels are labeled as individual cells and which pixels are labeled as background or boundary. This aligns with the nature of a dataset that contains many instances of dynamic *Bacillaria* colonies, including changes in movement phase, spatial orientation, and image opacity (e.g. the presence of algae that might mask the cells). For the Deeplab analysis, all images are rectified to a common orientation and hand-labeled with information about the desired properties of bounding box edges. We then obtain XML records for each image and randomly partition these data into two subsets: a training set (train.csv) and a testing set (test.csv). These files are used to generate a record file (TFRecord) which is used to move data in and out of the deep network.

More generally, Deep Learning is an instance of the neural network approach and relies upon a network with a large number of hidden layers relative to a standard neural network. We discuss neural networks in more detail in a previous section of this paper. In general, a deeper network with more hidden layers translates into a larger feature space. This provides a user with models that have better resolution, but also models that have a greater potential for error [50]. To promote reproducibility, a tutorial for DeepLabv3 implementation is available on Github [51], and the software implementation is available on Github (https://github.com/devoworm/Digital-Bacillaria).

#### Bounding Box Method (DeepLab v3)

Now we turn to the segmentation and measuring of individual *Bacillaria* cells. The first step is to employ methods for identifying the boundaries of cells, in particular, to distinguish the boundaries from the area of other cells and the image background. This is done by defining a bounding box and then applying a cell segmentation method to a single image (Figure 4). We define five points on each cell: a) the two ends of the cell (1 and 2 in Figure 4), b) the midpoint of the line formed between points 1 and 2 (3 in Figure 4), and c) the edges of a cell (4 and 5 in Figure 4) defined by drawing a line perpendicular to the axis defined by points 1, 2, and 3. Centroids are then calculated from these data in a post-processing step by finding the midpoints between and maximum values for the *x* (centroid *x*) and *y* (centroid *y*) axes.

**Figure 4.**
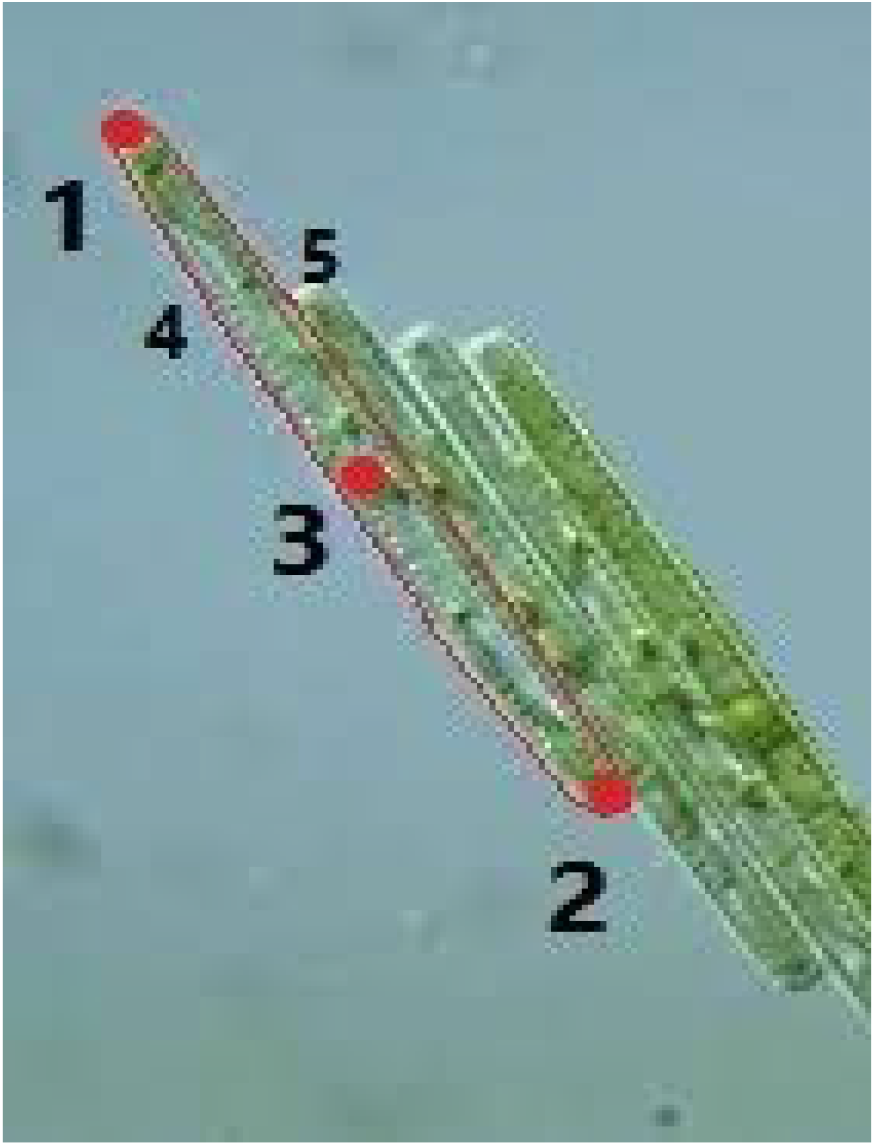
A diagram showing the five points on a sample cell (two ends, midpoint of the transverse line, and edges of the cell)

#### Soft Bounding Box Method (OpenDevoCell)

The OpenDevoCell platform was originally trained using pixel labeling and segmentation masks. Pixel level labeling requires a lot of effort and time, and does not always provide great results. A lot of data that we have is partially labeled, where instead of each pixel is given a zero or one label each pixel has a membership in a macro category. The data provides spatial information about these categories in the form of (*x*, *y*, *z*) coordinates or (*r*, θ) (polar coordinates). One way of approaching the problem is by broadly defining potential features in the form of boxes and refining the boxes using a deep Convolutional Neural Network (CNN) [52] to get a semantically segmented image where the segmented features are defined by distinguishing between specific labels.

The OpenDevoCell technique uses region proposal methods to generate segmentation masks. The candidate segments are used to update the deep CNN. The semantic features learned by the network are used to generate better candidates and proceed as an iterated procedure. This method was originally applied to *Caenorhabditis elegans* embryogenesis data by utilizing the spatial locations and making the bounding boxes. With ground-truthed bounding boxes, we can find the candidate masks that overlap the most with the bounding boxes. An error/cost function is used to maximize the overlap. For application to *Bacillaria* colonies, we convert our dataset to skeletons using a procedure implemented in GIMP 2.10. These skeletons are pre-masks that mimic the initial condition of the *Caenorhabditis elegans* embryos, which was a series of high-resolution images in which membrane expression of a GFP marker was used to define cell boundaries.

#### Noise Reduction (DeepLab v3)

Since edge detection is susceptible to noise in the image [53], the DeepLab v3 pre-trained model uses a noise reduction algorithm. The first step is to remove the noise in the image with a 5×5 Gaussian kernel which results in spatial smoothing [54]. To find the intensity gradient of a given image, smoothed (locally averaged) images are then filtered with a Sobel kernel [55] in both horizontal and vertical directions to get the first derivatives in the horizontal direction (*G_x_*) and vertical direction (*G_y_*). From these two images, we can find an edge gradient and direction for each pixel as follows:

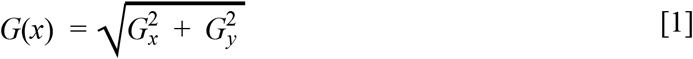

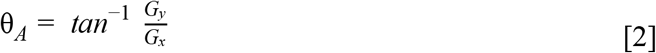

The gradient direction is always perpendicular to edges. It is rounded to one of four angles representing vertical, horizontal and two diagonal directions.

#### Non-maximum Suppression (DeepLabv3)

The DeepLab v3 pre-trained model also utilizes non-maximum suppression. After getting gradient magnitude and direction, a full scan of an image is done to remove any unwanted pixels which may not constitute an edge. For this, at every pixel, a pixel is checked if it is a local maximum in its neighborhood in the direction of the algorithmic gradient (see Figure 5).

**Figure 5.**
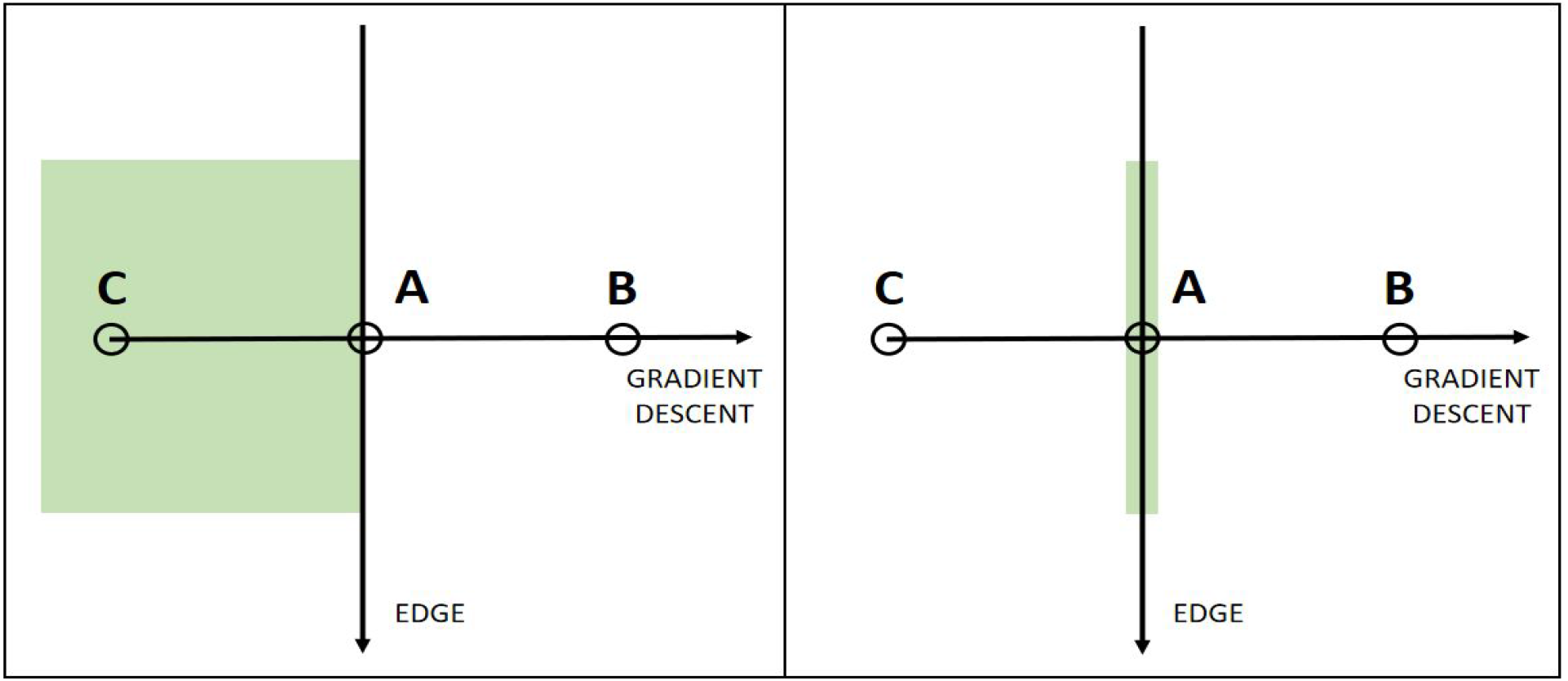
Point A is on the edge (vertical direction). The gradient direction is normal to the edge. Point B and C are in gradient directions. So point A is checked with point B and C to see if it forms a local maximum. If so, it is considered for the next stage, otherwise, it is suppressed (set to zero). The result is a binary image with “thin edges”

#### Hysteresis Thresholding (DeepLabv3)

DeepLab v3 also uses hysteresis thresholding [56] for image segmentation (Figure 6). Hysteresis refers to the retention of low threshold edges that are associated with high threshold edges. This stage of processing decides which edges in the image are most likely to be actual edges. Hysteresis thresholding relies on two threshold values, *V_min_* and *V_max_*. Any edges with intensity gradient more than *V_max_* are sure to be edges, while those below *V_min_* are sure to be non-edges and thus discarded. Those that lie between these two thresholds are classified edges or non-edges based on their connectivity. Hysteresis occurs when the number of edges identified by this method are fewer than those defined by V_min_, but greater than those defined by V_max_.

**Figure 6.**
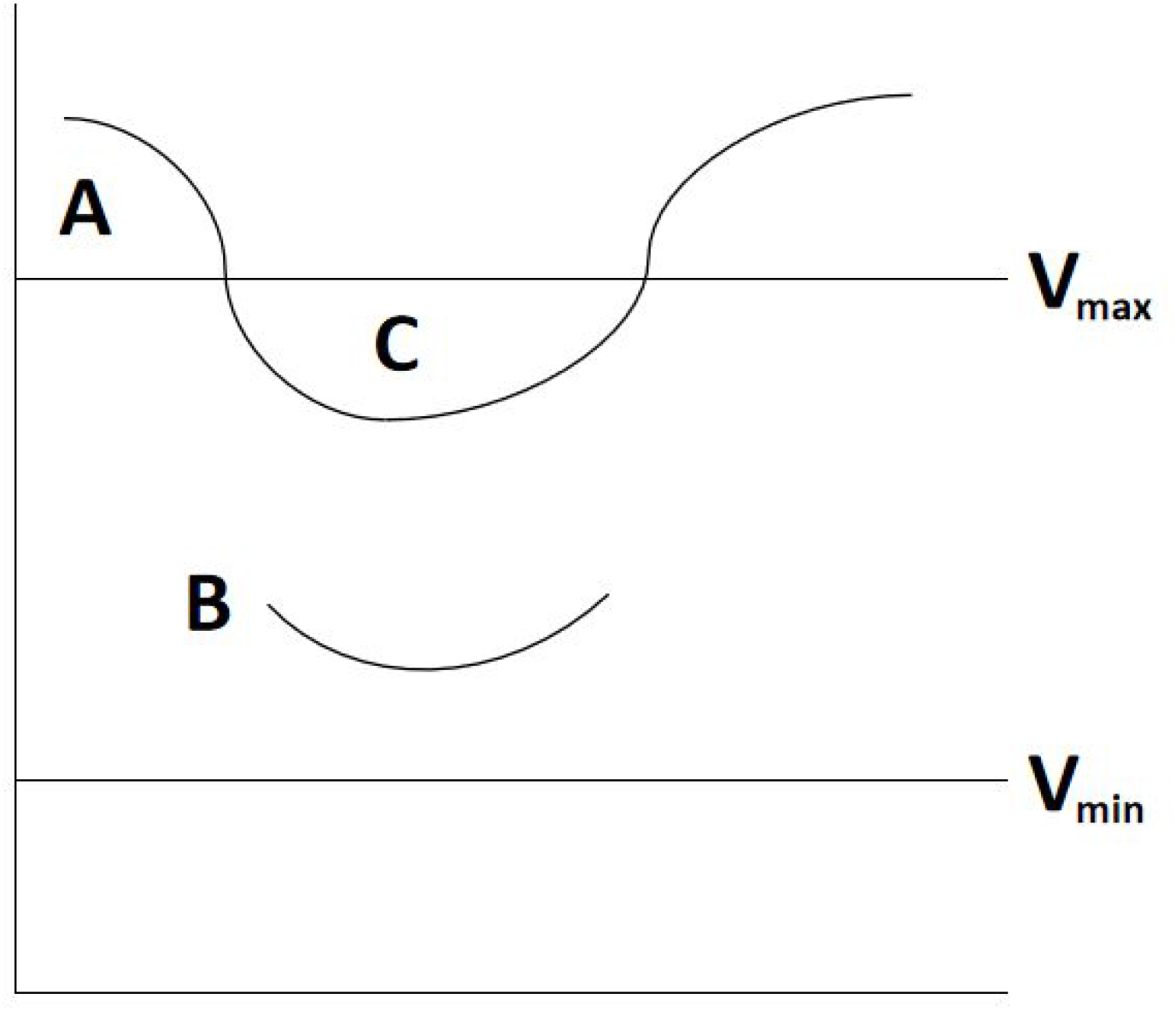
Diagram showing an example of hysteresis thresholding and the labeled edge relative to the “sure edge” threshold (V_max_)

The edge A is above the *V*_*max*_, so considered as “sure-edge”. Although edge C is below *V_max_*, it is connected to edge A, so that is also considered as a valid edge and we get that full curve. But edge B, although it is above minVal and is in the same region as that of edge C, it is not connected to any “sure-edge”, so it is discarded (Figure 6). It is very important that we have to select V_min_ and V_max_ accordingly to get the correct result. This stage also removes small pixels noises on the assumption that edges are long lines and ultimately produces strong edges in the image. Although this is quite an advanced technique, we are still not able to detect the cells when a colony stretches out during its course of the movement (see examples in Figure 1).

### DeepLabv3 Analysis

A number of methods were considered in the course of segmenting images for analysis by the DeepLabv3 model. The results for two of these (watershed and canny edge detection) are presented and contrasted here. The watershed and Canny edge detection analyses are done in OpenCV (Intel Corporation, Santa Clara, CA). Two methods are presented as a means of comparing performance on static images, and then these methods are contrasted with the deep learning results.

#### DeepLabv3 Model Dataset

Our dataset used as input to the pre-trained model has been extracted from YouTube videos of *Bacillaria* colonies. The image backgrounds are normalized and rotated to be in horizontal alignment. The data are normalized using a *z*-score transform (or *y*-score transform for images with an n < 6). This creates a coordinate space that is based on individual colonies relative to the mean and standard deviation of the full dataset. The full dataset of images and segmented cells (*N* = 65, *n* = 810) is also paired down to a dataset of selected samples (*N* = 46, *n* = 599). The selected dataset is also analyzed using a Principle Component Analysis (PCA) using SciLab 6.0 [57].

#### Feature detection methods

The Watershed method [58] is based on the concept of geographic watersheds, or drainage basins. The relative contrast of pixels across the image is used to define the watersheds (high-intensity regions) and the boundaries between watersheds (low-intensity regions). This is done by treating image intensities for each pixel as part of an elevation map [59], which results in a binary classification of the image. The Watershed algorithm is particularly good at finding contours between distinct regions of an image. Thresholding of overall image intensity was done using a marker-based approach (optimization via trial-and-error).

Canny edge detection [60] operates on a noise-filtered intensity gradient derived from the original image data. As with the watershed method, Canny edge relies on a series of thresholding and filtering techniques to determine the strength of potential edges in the image. One of these is to rely on averaging and signal suppression to classify all potential edges into a set of four angular orientations (0, 45, 90, 135) across a 180-degree arc. Canny edge detection also involves a hysteresis step, which is employed to deal with ambiguous components of the signal relative to an intensity threshold [61].

#### Primary Dataset

We collected microscopy movies of movement for a single *Bacillaria* colony using light microscopy. These data were converted into still images, which are transformed and segmented using machine learning techniques. Microscopy is conducted using a Zeiss standard upright microscope. All images are brightfield, 40x plain objective. Images are extracted from videos representing 8x time-lapse. Data are collected from a colony transferred to a slide from culture.

### Primary Dataset Analysis

We will also present an analysis of a primary dataset. Part of this involves masking and segmentation of microscopy images using the OpenDevoCell platform. The other part of this analysis involves using a time-lapse approach to track the motion of cells over time. We are able to approximate patterns of movement and acceleration by identifying labeled features across images over time.

#### OpenDevoCell

We analyzed the primary data using the OpenDevoCell application (https://open-devo-cell.herokuapp.com/). For this analysis, there is a masking step and a segmentation step. OpenDevoCell is a Java-based deep learning platform that uses the TensorFlow library Deep Learning 4J.

#### Time-lapse analysis

We also analyzed the primary data using time-lapse techniques. The time-lapse is created using VirtualDub, version 1.10.4.35491 (http://www.virtualdub.org/). The *Bacillaria* strains contained in our primary data (videos) have been harvested from the Neckar river in Germany (49°04’41.8”N 9°09’17.9”E). Samples were collected on September 14, 2019. The average size of each cell (filament) is approximately 81µm.

### Data Availability

Select unprocessed (raw) data are available at our Github repository (https://github.com/devoworm/Digital-Bacillaria), processed numeric and image data (numeric tables and skeletonized images), and select video files are available on the Open Science Framework (DOI 10.17605/OSF.IO/AR8C3).

## Results

### Watershed segmentation and Canny Edge Detection

Neither the Watershed Segmentation nor the Canny Edge Detection approaches provide very strong performance. The desired feature (closed boundary around each cell) was not detected, as interference from noise in the form of other algae or unclear cell boundaries dominates the analysis. Examples of these results are shown in Figure 7.

**Figure 7.**
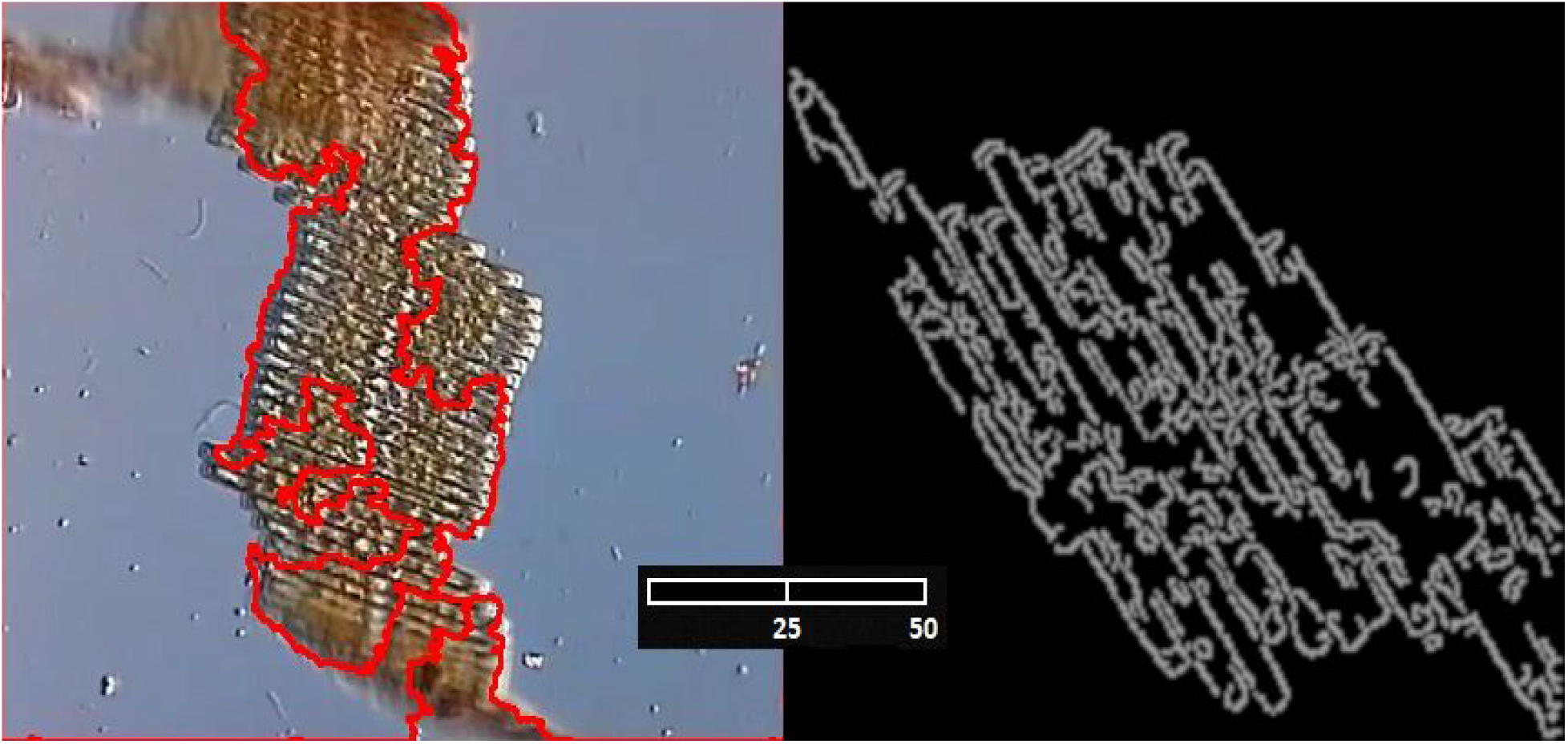
An example of feature identification performance for the Watershed Segmentation algorithm (left, red boundary) and Canny Edge detection algorithm (right, white boundary). Image scale: 38.36 μm per cm, or 0.325 μm per pixel. Scale bars 50 μm

### Deep Learning

The results for the pre-trained model (DeepLab v3) were much more accurate. Based on feature training using the bounding box method demonstrated in Figure 8, we are able to reconstruct several parameters of the individual cells which suggest an accurate reconstruction of the source image. We can also see the improved accuracy of the correct performance in Figure 9. If the source image is not too blurry (out of focus, dominated by artifacts), the model can be trained and features detected without too much difficulty.

**Figure 8.**
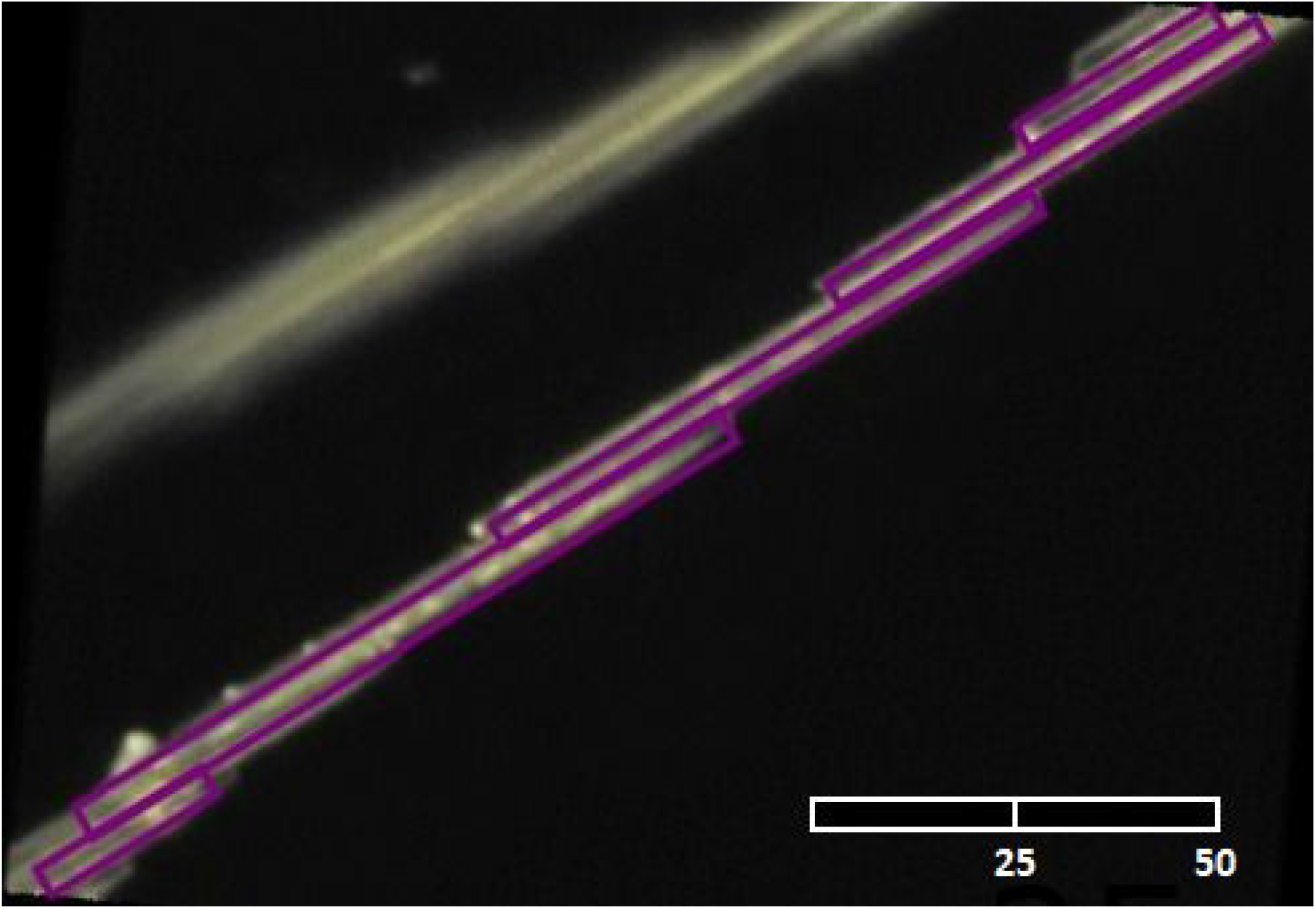
An example of feature identification training (purple rectangles) for the deep learning approach on a single set of cells. Notice the resolution of the colony. An example of correct performance is shown in Figure 5. Image scale: 38.36 μm per cm, or 0.325 μm per pixel. Scale bar 50 μm

**Figure 9.**
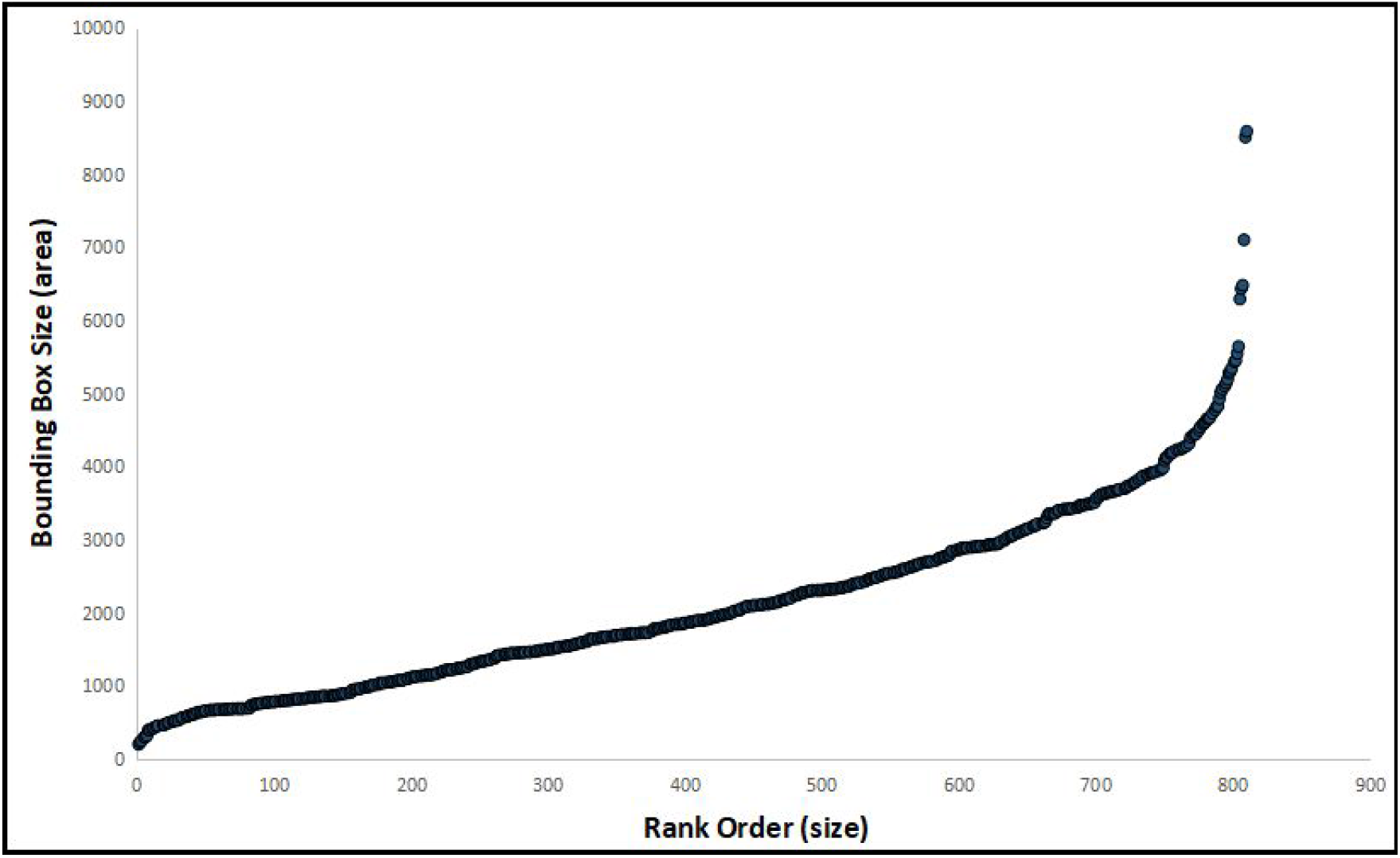
Rank-order analysis of bounding box (cell) sizes (area) across the dataset. The area is measured in pixels squared. Image scale: 38.36 μm per cm, or 0.325 μm per pixel

An analysis of the pre-trained model outputs is shown in Figures 9, 10, and 11. Figure 9 shows the distribution of cell sizes across the full dataset, ranked from smallest to largest. There is a long tail of very large cells (750 to 800 pixels^2^) that represents filaments much larger than their neighbors. This could be due to errors in the segmentation process of bounding boxes, but could also represent two cells lumped as one, or cell division in the process. Figure 10 also uses a rank ordering from largest to smallest instance but breaks Figure 9 down into the lengths and widths of each bounding box. In this graph, we see that the length is generally larger than the width as expected, but that there is a more normal distribution of bounding box lengths. This could mean that there is natural variation in the length, but this could also be partially due to resolution loss across the image or the truncation of cells at the edge of the image.

**Figure 10.**
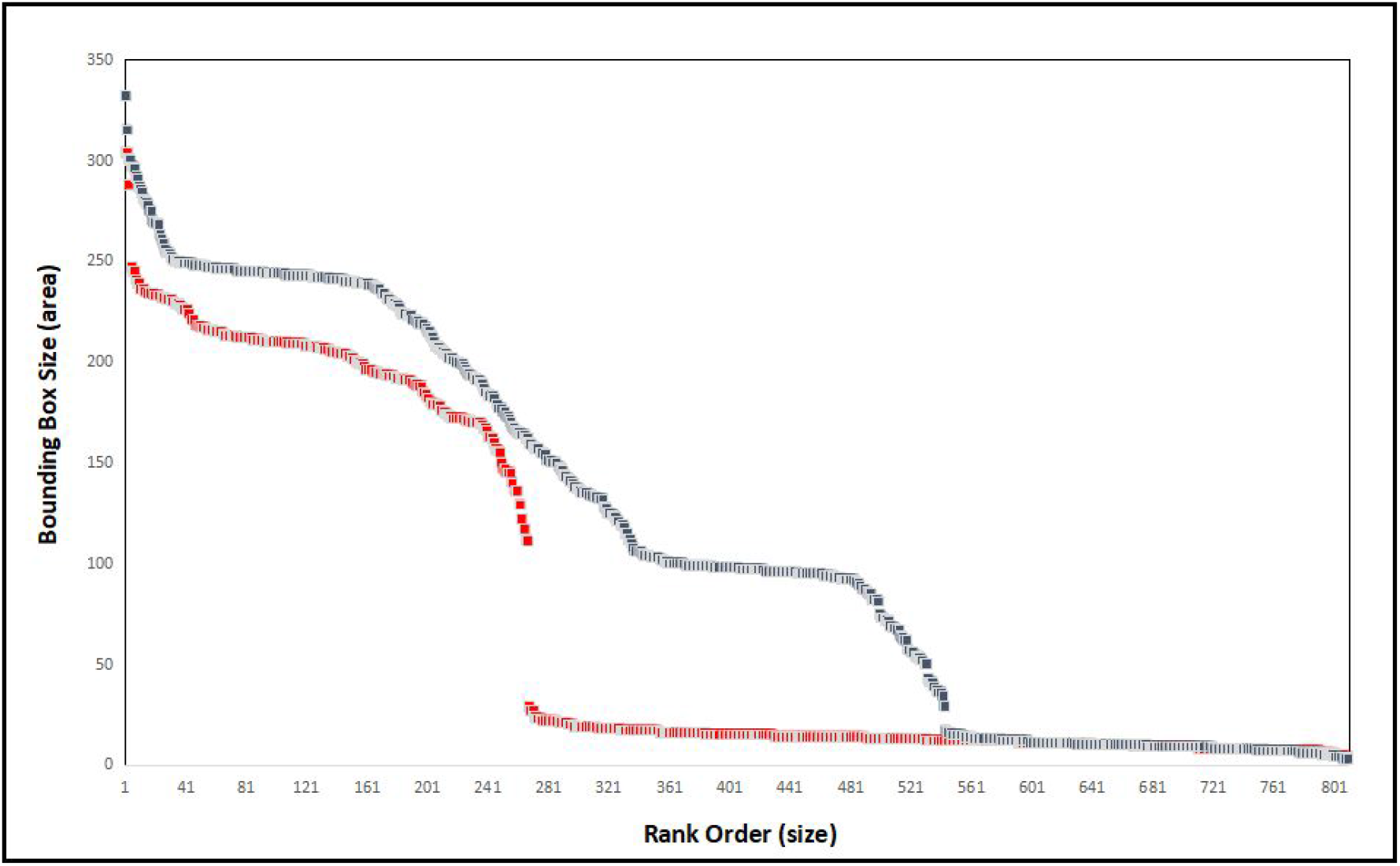
Rank-order analysis of height (blue) and width (red) of bounding boxes (cells) across the dataset. The area is measured in pixels squared. Image scale: 38.36 μm per cm, or 0.325 μm per pixel

**Figure 11.**
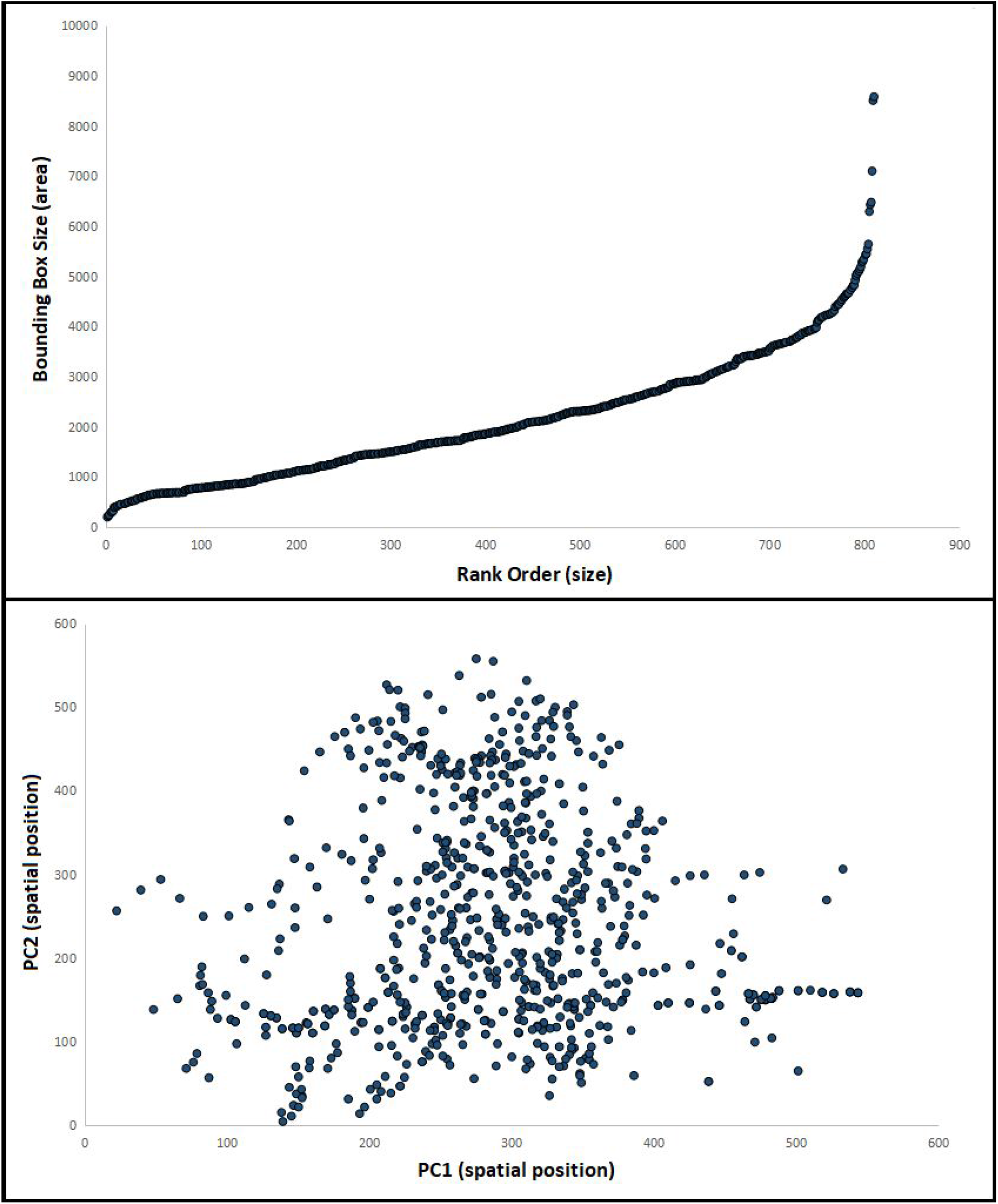
Top: location of centroids in normalized coordinate space in selected dataset for static analysis. Bottom: First two principal components from PCA analysis (PC1 represents horizontal position, while PC2 represents vertical position) of coordinates representing the *x,y* position for all four edges of each bounding box using the selected datasets for static analysis. Image scale: 38.36 μm per cm, or 0.325 μm per pixel

Figure 11 contains two graphs representing different aspects of the selected dataset. The first relationship (top) is a bivariate plot of all centroid locations among the selected data, similar to the rank-order analysis of bounding box areas shown in Figure 9. In the second relationship (bottom). A Principle Component Analysis (PCA) is conducted and represented by plotting the first two principal components. Both plots use a normalized coordinate system (see Methods).

Figure 12 shows how the pre-trained model can be optimized. In this case, we tried two different optimizations to extract the bounding boxes from random images in the training set. Each optimization (as well as the final segmentation) provides a tradeoff between the number of boxes and the accuracy of the segmentation set. The final segmentation included elements of each optimization, which is a tradeoff between a greater number of boxes and an increase in false positives. Even the most optimized result will produce false positives and false negatives due to differences in the resolution of cell boundaries across the image.

**Figure 12.**
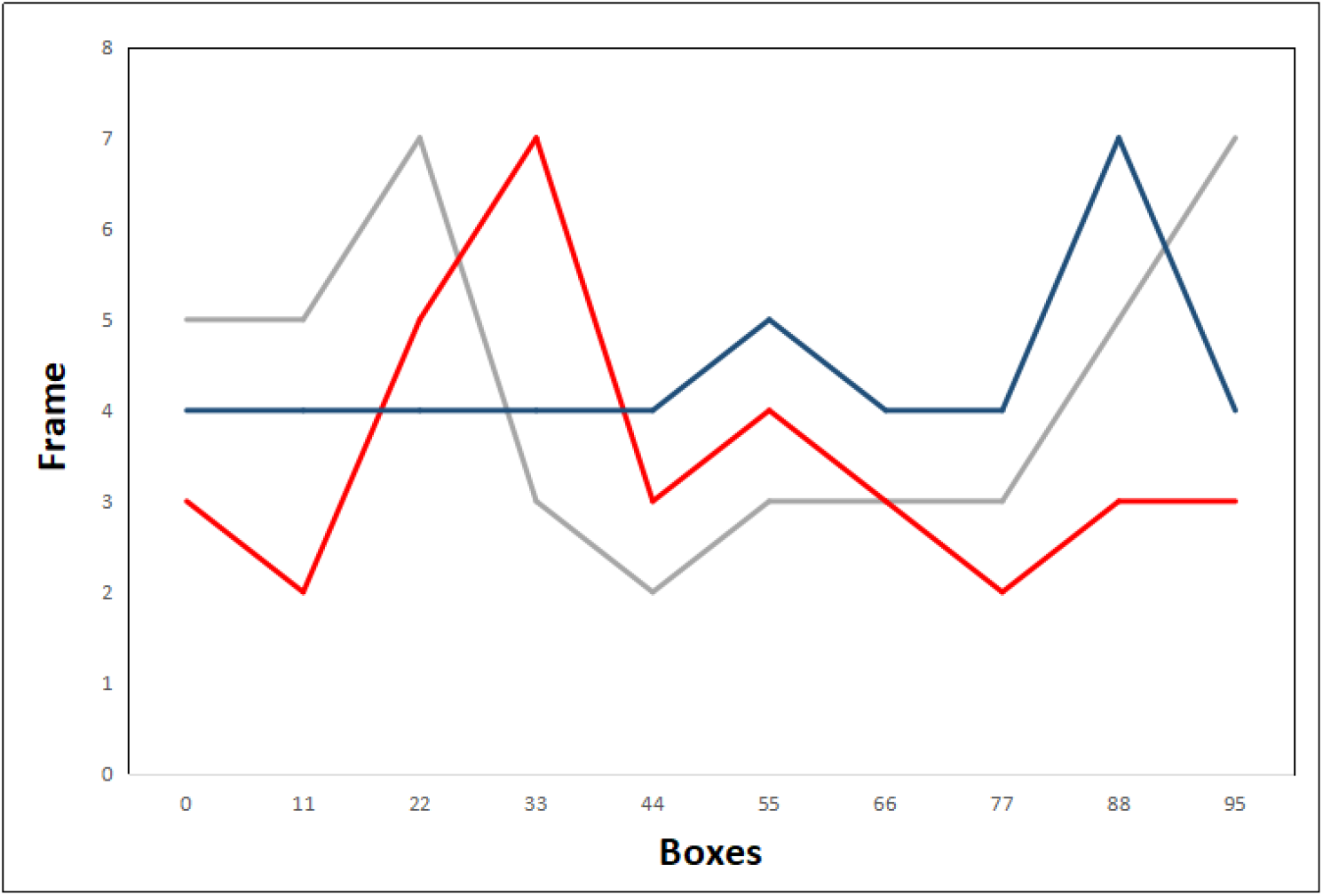
An example of feature identification optimization procedures implemented in DeepLab v3. GRAY: no optimization applied, RED: Optimization #1, BLUE: Optimization #2. Given an initial number of training frames (y-axis), the non-optimized procedure (originally detected) will yield a certain number of boxes (x-axis). Applying various optimization procedures generally leads to a decreased number of boxes per frame for both low and high numbers of boxes

Although this optimal tradeoff between feature number and accuracy was used in our analysis, Figure 13 demonstrates how difficult it is to achieve a perfectly accurate segmentation. In Figure 13, four examples of segmented images are shown with centroids of the segmented cells plotted in a bivariate coordinate space on the left and the original image on the right. For images A, C, E, and G a normalized coordinate space (see Methods) is used for both the x and y axes. While it is hard to see the degree of concurrency, there is a correspondence between the centroid locations and the cells as stacked in their respective colonies. In general, images such as A and C are harder to resolve than images such as E and G. This is likely due to the configuration of the colony as a “V” (as is the case with E and G) shape versus a more irregular shape.

**Figure 13.**
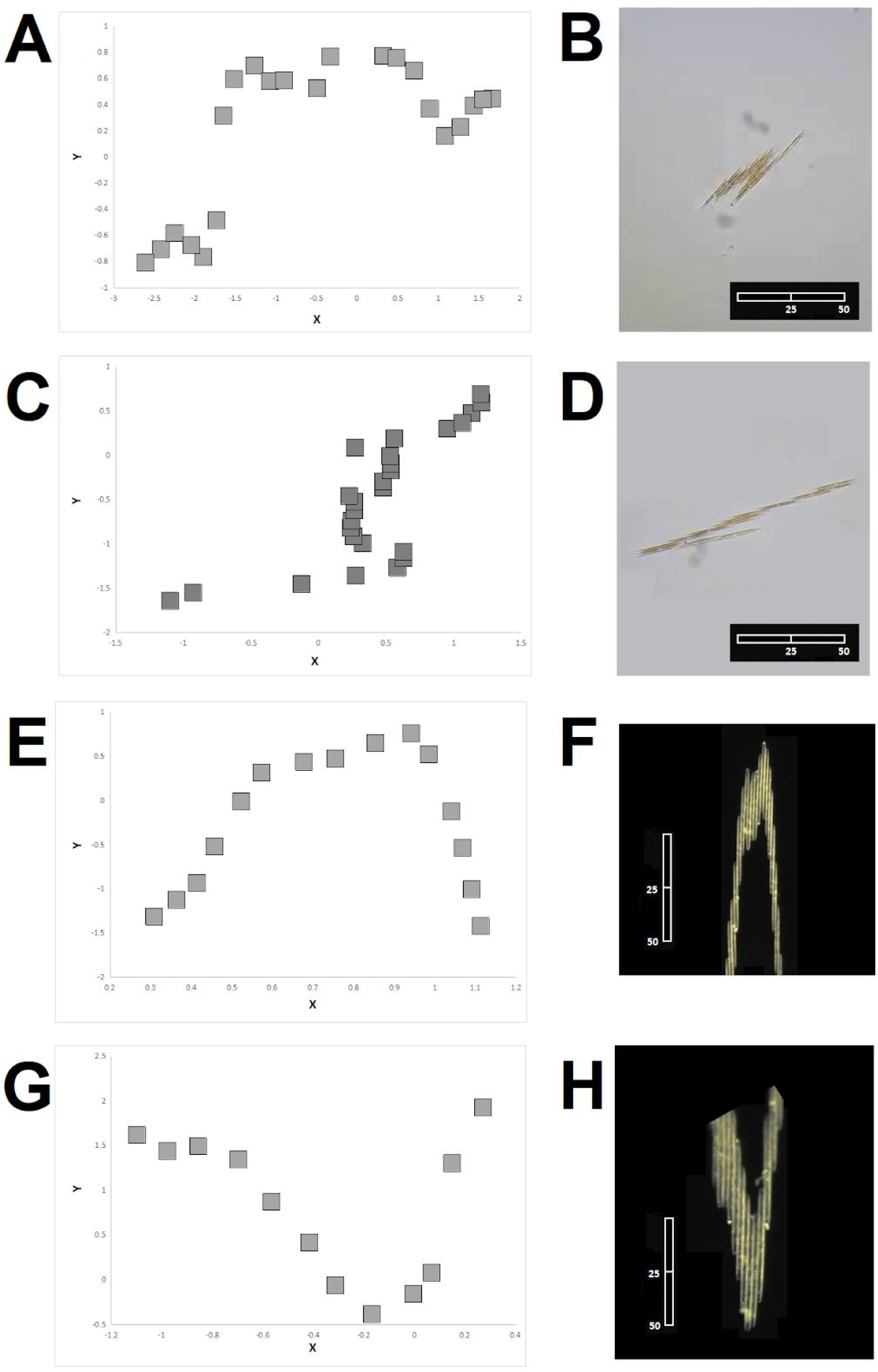
Four examples of how the identified features map to two different images (A, C, E, G) of a *Bacillaria* colony. Points (B, D, F, H) represent the centroids for all bounding boxes identified in images A, C, E., and G, respectively. Image scale: 38.36 μm per cm, or 0.325 μm per pixel. Image scale: 38.36 μm per cm, or 0.325 μm per pixel. Scale bars 50 μm

Now we will use the OpenDevoCell model to demonstrate what happens when a model trained for one type of biological system (nematode embryos) is utilized to identify cells in another context (*Bacillaria* colonies). This analysis does not demonstrate the efficacy of the OpenDevoCell model, rather, it is to show how we might go beyond the bounding box method to more general implementations. Generally, models that learn from data are generalized to a specific set of features. Our purpose here is to see if there are any similarities between nematode embryogenesis and diatom functional morphology. The first step in this demonstration is to create skeletons of the original images in order to mimic the resolution of a suitable image for OpenDevoCell, which has been trained to recognize GFP-labeled cell membrane boundaries (between cells). We show the original images along with their corresponding image skeletons in Figure 13 using three exemplar images from the primary data described in the Methods section.

The pre-masking exercise shown in Figure 14 demonstrates how difficult it is to define to distinguish an outer edge versus the contours and internal features of *Bacillaria* cells. Yet along with the templating analysis featured in Figure 15, we demonstrate that intracellular features can be identified and potentially used as quantitative features. Using the DeepLabv3 pre-trained model, we can derive bounding boxes with significant variability in how they map to the original image. In this case, we create image skeletons that are interpretable by the OpenDevoCell model. As the OpenDevoCell model has been optimized for fluorescent images, these image skeletons are bright green and can be separated from both the background and extraneous noise. When these skeletons are presented to the OpenDevoCell model, it can provide spatially-referenced segmentation of *Bacillaria* colonies. These objects can also be co-registered with bounding boxes yielded from the DeepLab analysis.

**Figure 14.**
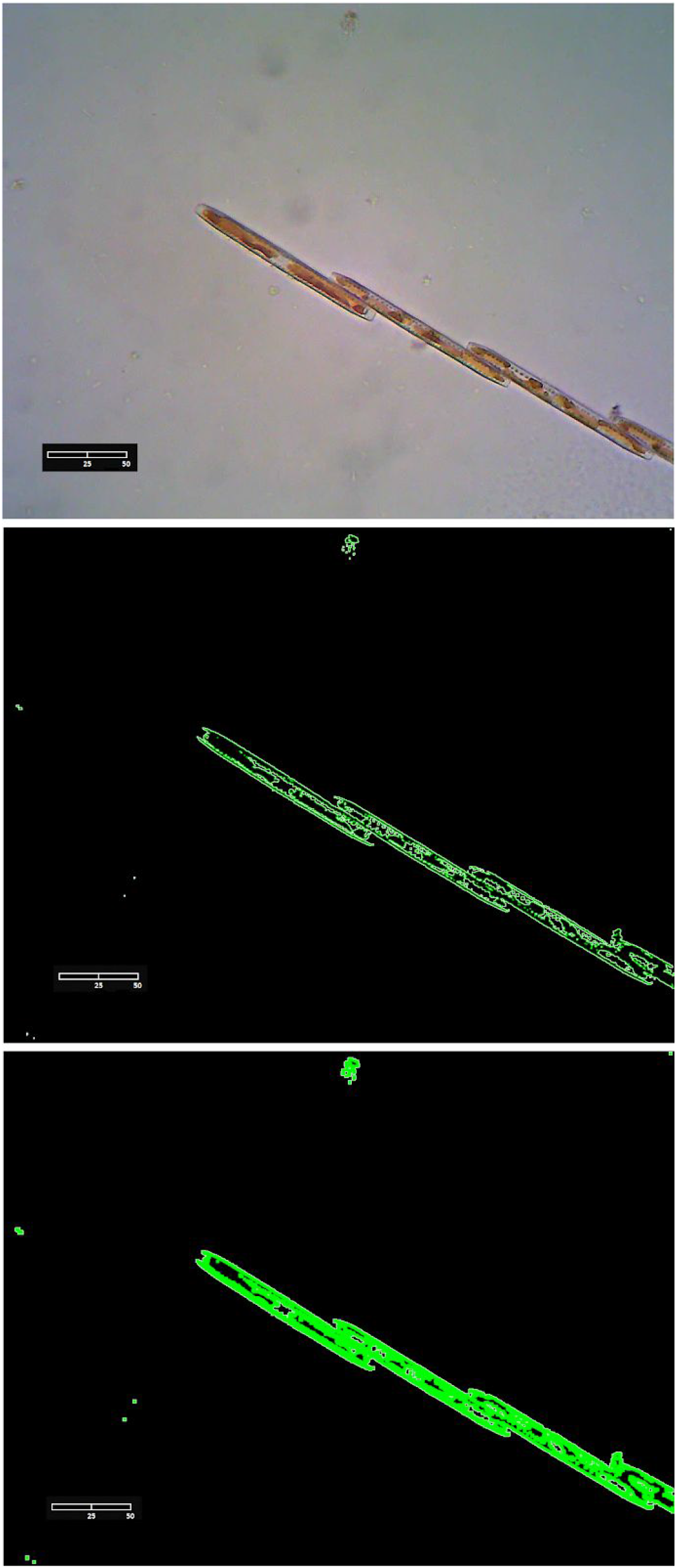
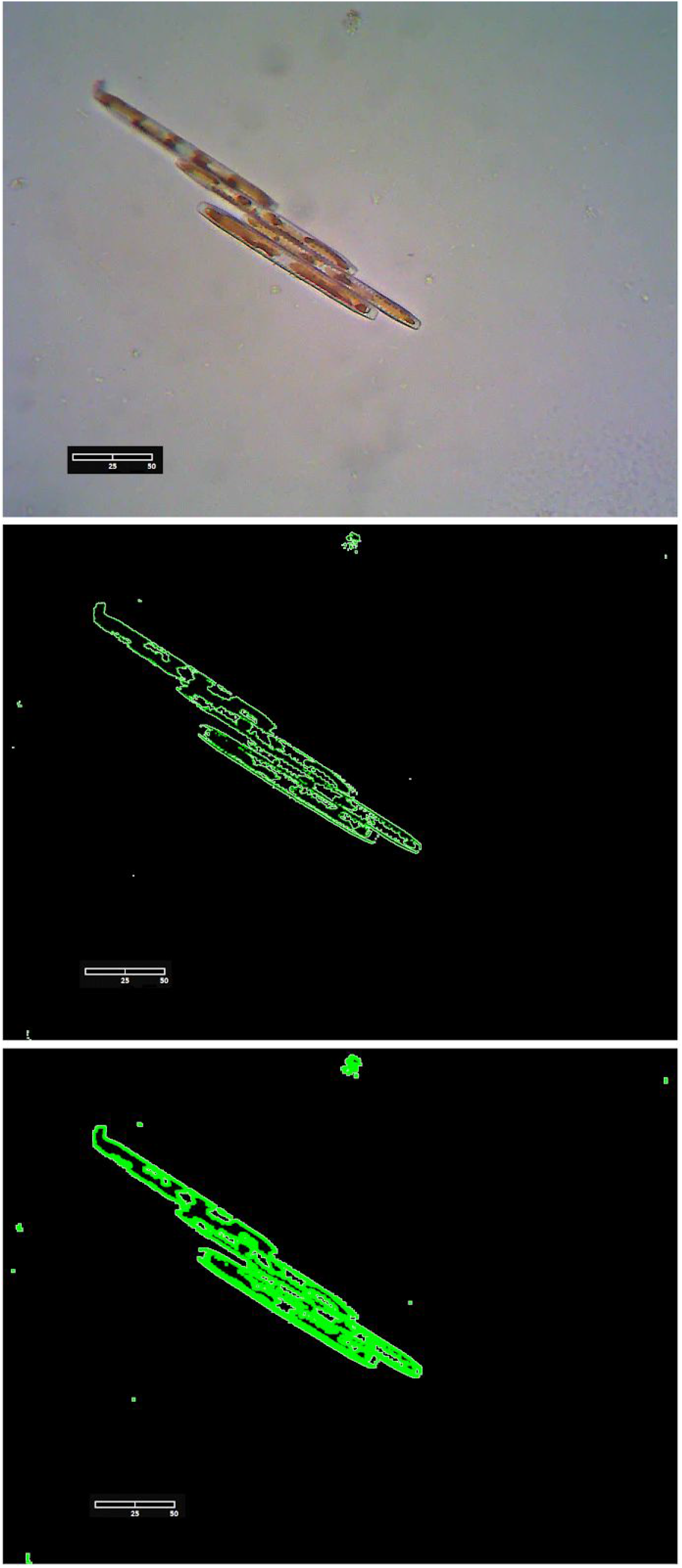
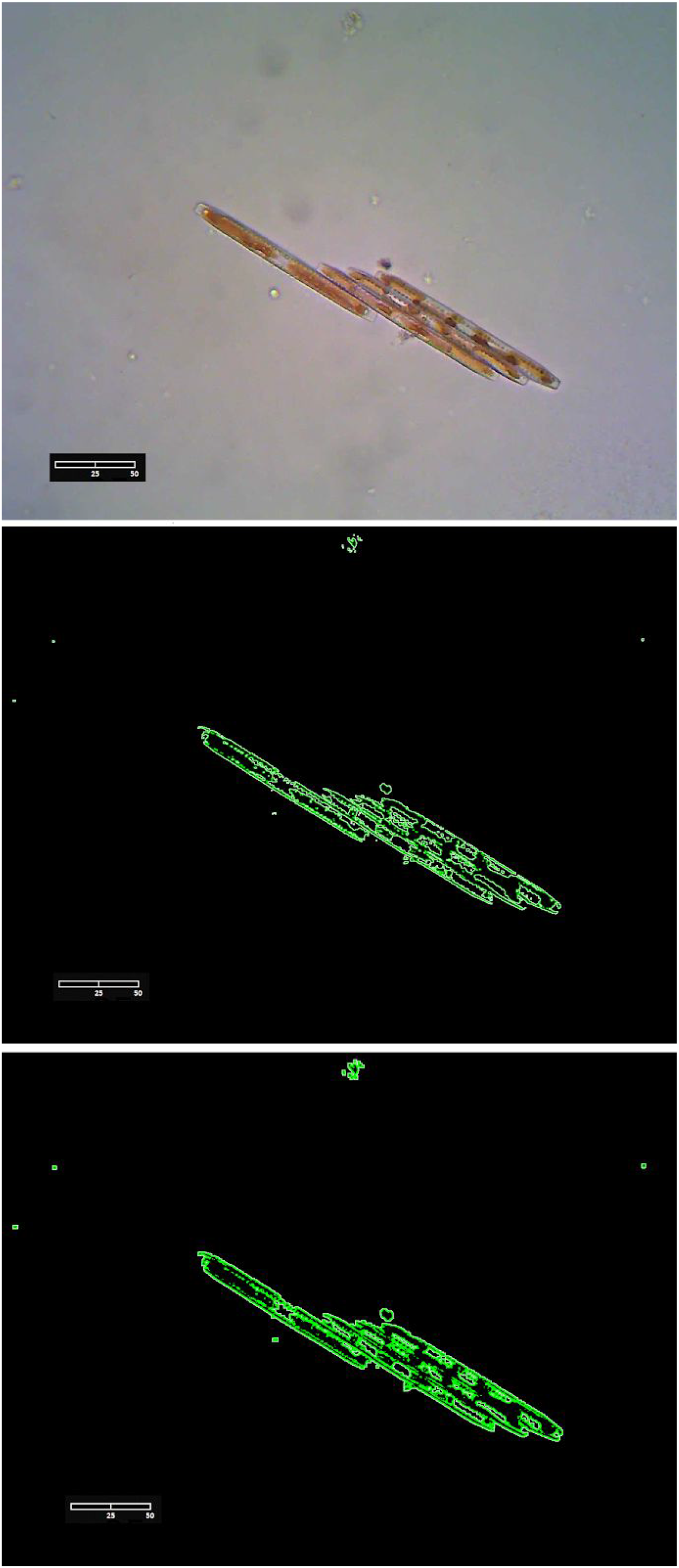
Three examples of how images of a *Bacillaria* colony are converted into a skeleton image. TOP ROW: light microscopy images, MIDDLE ROW: thin skeletonization based on a procedure implemented in GIMP, BOTTOM ROW: thick skeleton based on a procedure implemented in GIMP. Image scale: 38.36 μm per cm, or 0.325 μm per pixel. Image scale: 38.36 μm per cm, or 0.325 μm per pixel. Scale bars 50 μm

**Figure 15.**
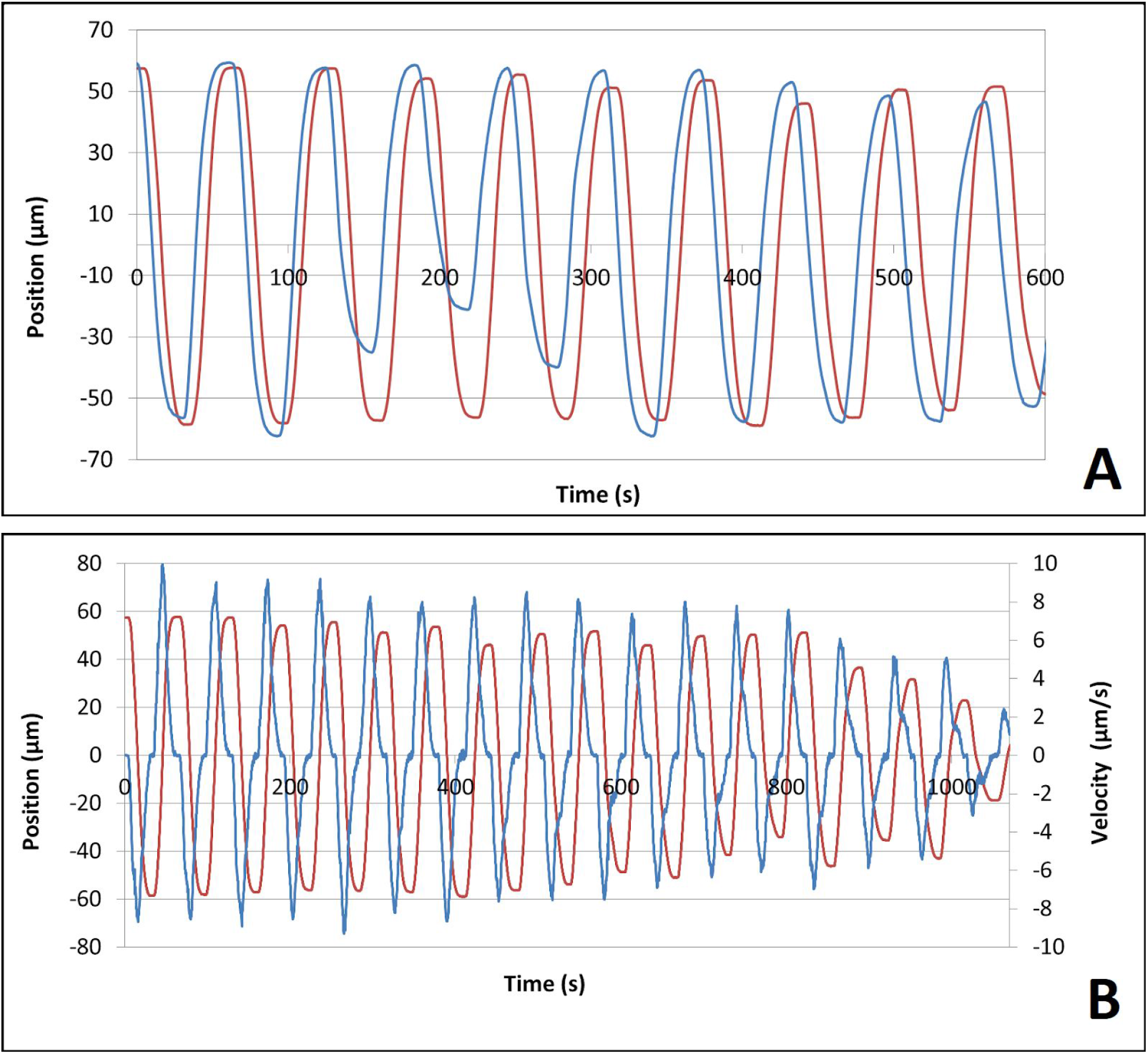
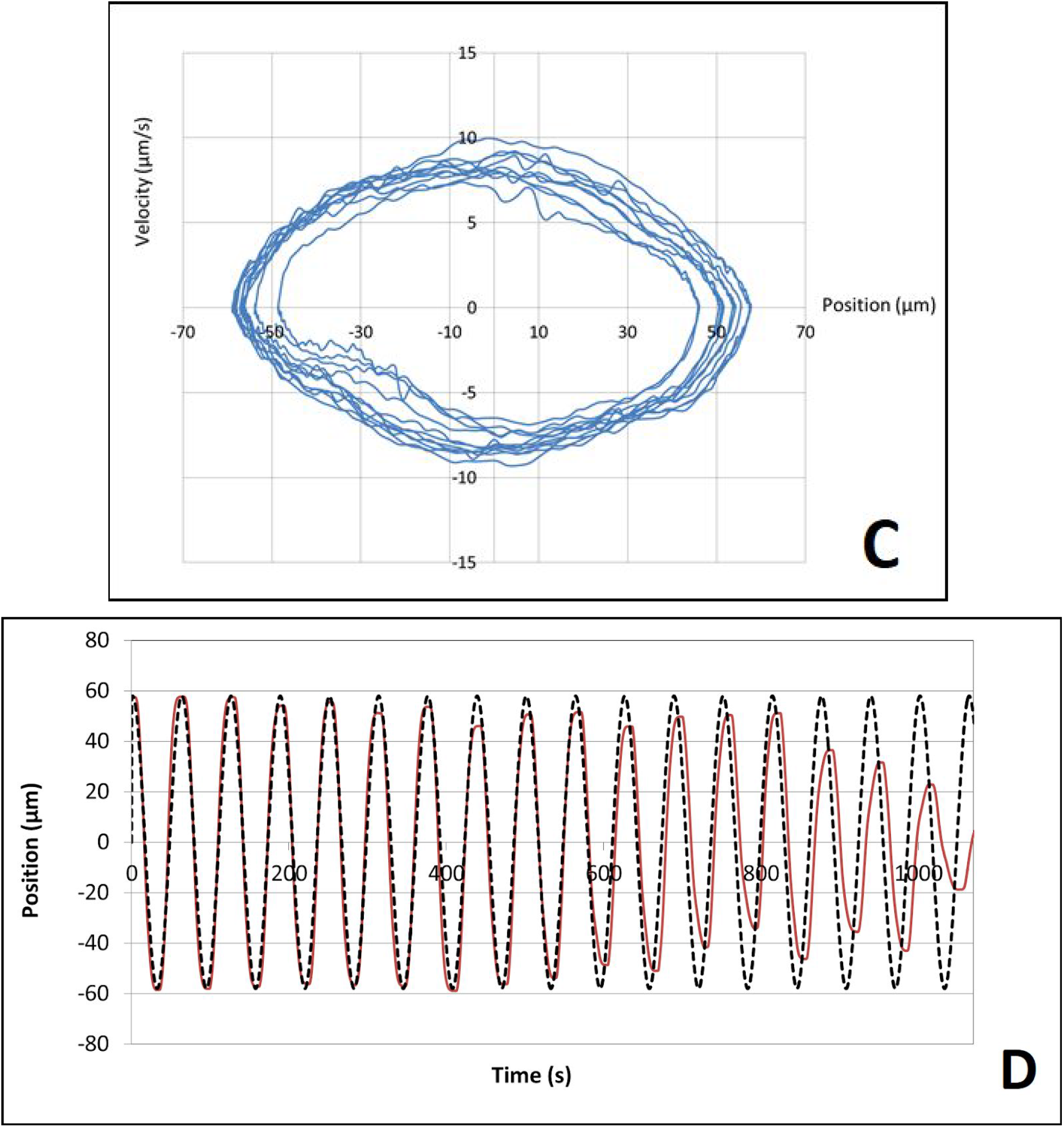
Examples of relative movement of cells in a sample colony. A: Comparisons between changes of position for cell #2 relative to cell #1 (red) and cell #3 relative to cell #2 (blue). B: Comparisons between changes of position (red) and changes of velocity (blue) for cell #2 relative to cell #1. C: a phase diagram of the data shown in B, velocity versus position. D: comparison between changes of position for cell #2 relative to cell #1 (red) and sine wave (black dashed). The oscillation period in frame C is 62.56 seconds

In this case, a model that works quite well for embryos performs more like the unsupervised models shown in Figure 7. One reason why the DeepLab model may be more successful at defining bounding boxes in this context could be the ability to recognize generic rectangles, features which are helpful for identification but not descriptive of subtle changes in structure or shape. This is particularly relevant in terms of cells that fall partially out of frame or detail within what is captured as a bounding box. While difficult to obtain, the objects segmented by this implementation of OpenDevoCell are more detailed and less square than those extracted by the DeepLab analysis. We propose that a combination of OpenDevoCell and pre-masking may be useful in revealing potential intracellular features. This is particularly true for capturing variations within cells, which allows for these features to be mapped to the bounding box segmentation data. This would provide a nice balance between accuracy and detail, but also yields inconsistent results for features that change position between frames over time.

Given our limitations in acquiring feature/background separation for sequential samples, our final analysis is to apply motion tracking to the *Bacillaria* colonies across frames. This can be done using template analysis (see Methods). The templating method shown in Figure 3 shares commonalities with the DeepLab model in how cells are segmented and represented. In both cases, the centroid approximation of an identified feature is used as a reference point. Yet for purposes of approximating motion over time, the most important information for the analysis of the movement is the positions of individual cells in a *Bacillaria* colony relative to neighboring diatoms as a function of time. Highlights of our analysis for single-cell motion over time, particularly with respect to neighboring cells, is shown in Figure 15.

At the level of individual cells, the movement of the colony appears to be oscillatory. Comparisons of cell #2 and cell #3 reveal oscillations that are slightly out-of-phase (Figure 15A). As we might expect, changes in velocity for the oscillation of a single cell are noisier than changes in position for that same cell (Figure 15B). Yet the velocity function shows a relatively smooth transition between two extremes. When the entire chain comes to rest, we observe a dampening of both the position and acceleration time-series. In Figure 15C, the absolute value of velocity increases linearly with time until the diatoms in consideration lie next to each other with their apices, then decreases linearly until they slow down. Linear increases and decreases of speed over time means that the curve of the positions is composed of parabolic segments. For this comparison of position and velocity, we are able to replicate the findings of Yamaoka et.al [8]. An exception to this are the ranges in the proximity of the reversal points, in which this linearity is not given. In our analysis, the diatoms behave as if velocity is proportional to the length of their common contact surface.

## Discussion

This paper introduces a new approach to understanding biological processes more generally and specifically *Bacillaria* colony morphology and movement. We employ image processing and machine learning techniques to segment images and extract quantitative parameters. These data can then be used to both infer the phenotypic structure of a colony and movement patterns of these colonies. Of particular interest is the combination of multiple image processing and deep learning techniques with a biomechanical analysis. Hopefully, this will provide guidance towards the future development of digital models.

This is the first attempt at characterizing individual colonies and cells using advanced computational techniques. Future refinements of analytical techniques and input data will allow us to build better models, bridging the gap between prior work and emerging methods. In addition, we provide our own innovations to the study of *Bacillaria*. While our approach provides unique information about the process of colony form and function, we are also in a position to develop and clarify potential theoretical arguments regarding organismal phenotypes, movement, and behavior.

Even when the various aspects of feature selection are optimized (as shown in Figure 12), false positives are occasionally included in the output. This is true for all the techniques presented here. For example, the segmentation process can introduce errors when the input data is highly variable. Despite our attempt to normalize both input and output data, the normalization procedures make certain assumptions about the data that do not fully capture natural variation. More generally, the shape of colonies observed in our input data does not represent every configuration found in nature. While we expect our analyses to be robust to variations on observed shapes and movement dynamics, we cannot assure that this is the case.

Manipulations such as tracking and segmentation of diatoms colonies work best when the colony lies in the focal plane perpendicular to the optical axis for a sufficiently long time. This is likely to be the case for both the tracking experiments and the secondary data. Since *Bacillaria* colonies often move in three dimensions, diatoms are often seen from different perspectives and overlaps may occur. Although it is possible to force the colonies between a slide and a cover glass into this position, the movement is often influenced by adhesion to the confining surfaces. We must also consider the role of hydrodynamic flow in distorting the movement function within and between measurements. Yet since *Bacillaria* lives at low Reynolds numbers, it is likely to have minimal impact on the results. In general, analysis is most accurate in cases where small colonies are in continuous contact with the visible surface.

These issues might be overcome with a larger and more diverse training set [39, 41]. The fact that the number of cells in a colony is constant from one frame to the next (unless a cell division has occurred) has not yet been incorporated. The lack of interpolation for cells that are truncated by the edge of the microscopy image frame is another issue. While this method is able to generate useful quantification of the image set, we still arrive at a number of issues with mapping the functional phenotype of a *Bacillaria* colony to a digital representation. This is a meta-issue when compared to false-positive identification, and so may require a new concept to characterize the relative imprecision of the mapping between image and digital representation. Better methodology with respect to the feature space (more subtle components of colony morphology) would also be a helpful future advance.

This paper also makes several contributions to both computational biology and machine learning literature. The first is to create a digital model of several parameters that are implemented by an existing general-purpose pre-trained model. This has largely been successful and has allowed us to extract several key parameters that describe the phenotype. We are also able to compare these results with models trained for other specific biological tasks. This was less successful but does guide us toward future work. This digital model will be made available as an open-access model and can be updated with improved data and methods. Another contribution is to create a dynamic model of the *Bacillaria* that will allow us to predict movement and other deformations of the phenotype. This will help us characterize not only modes of movement, but also any potential collective behaviors that require coordinated decision-making between cells.

A secondary theme of our paper are cases in which deep learning techniques succeed or fail at capturing the desired features. In the results section, unsupervised techniques are ruled out as inadequate, while the more technically sophisticated deep learning techniques are shown to yield results of varying utility. Of particular interest is the level of generalizability for such models. The optimal model should not be limited by biological variability, but should also be able to identify uniquely biological features, even when they mimic mechanical features. One solution to this are toy models, which can be used to capture complex processes in a simplified and mathematically-tractable model [62]. Results of the comparison between the DeepLabv3 and OpenDevoCell models suggest the need for a specialized pre-trained model [63] optimized for the shape, movement patterns, and intra-cellular contours of a *Bacillaria* colony (see Figure 2). Specialized pre-trained models have been created for a host of specific types of systems such as linguistic and object recognition and transfer, so creating a model specialized for the analysis of dynamic biological systems is both desirable and attainable.

## Acknowledgments

We would like to thank members of the DevoWorm group, in particular, the two working groups within DevoWorm (Digital Bacillaria and DevoWormML) that made this paper happen. Thanks also go to the diatom community for access to movies and other information on an otherwise obscure organism, and open-source software community for developing the tools used in the presented analyses. Special thanks go to the Google Summer of Code program for funding and enabling a portion of this work. Additional thanks go to Matt Ashworth for providing additional *Bacillaria paradoxa* specimens from Florida, USA.

## Notes

https://osf.io/ar8c3/

https://github.com/devoworm/Digital-Bacillaria

### Competing Interest Statement

The authors have declared no competing interest.

### Summary of Updates

This version reflects revisions that correct minor mistakes, add additional references, and clarify a few sections of the text (including figures).

